# Novel therapeutic strategies for injured endometrium: Autologous intrauterine transplantation of menstrual blood-derived cells from infertile patients

**DOI:** 10.1101/2022.11.17.516854

**Authors:** Satoshi Hosoya, Ryo Yokomizo, Harue Kishigami, Yukiko Fujiki, Erika Kaneko, Mitsuyoshi Amita, Takakazu Saito, Hiroshi Kishi, Haruhiko Sago, Aikou Okamoto, Akihiro Umezawa

## Abstract

**Background:** Menstrual blood-derived cells show regenerative potential as a mesenchymal stem cell and may therefore be a novel stem cell source of treatment for refractory infertility with injured endometrium. However, there have been few pre-clinical studies using cells from infertile patients, which needs to be addressed before establishing an autologous transplantation. Herein, we aimed to investigate the therapeutic capacity of menstrual blood-derived cells from infertile patients on endometrial infertility.

**Methods:** We collected menstrual blood-derived cells from volunteers and infertile patients, and confirmed their mesenchymal stem cell phenotype by flowcytometry and induction of tri-lineage differentiation. We compared the proliferative and paracrine capacities of these cells. Furthermore, we also investigated the regenerative potential and safety concerns of the intrauterine transplantation of infertile patient-derived cells using a mouse model with mechanically injured endometrium.

**Results:** Menstrual blood-derived cells from both infertile patients and volunteers showed phenotypic characteristics of mesenchymal stem cells. *In vitro* proliferative and paracrine capacities for wound healing and angiogenesis were equal for both samples. Furthermore, the transplantation of infertile patient-derived cells into uterine horns of the mouse model ameliorated endometrial thickness, prevented fibrosis and improved fertility outcomes without any apparent complications.

**Conclusions:** In our preclinical study, intrauterine transplantation of menstrual blood-derived cells may be a novel and attractive stem cell source for the curative and prophylactic therapy for injured endometrium. Further studies will be warranted for future clinical application.

## Background

Infertility is defined as failed conception after 12 months of regular unprotected sexual intercourse and it is estimated to affect 8-12% of couples of reproductive age ^1^. The number of worldwide infertile patients has increased to more than 186 million people, the majority being residents of developed countries ^2^. Among various etiologies of infertility, endometrium function is an essential factor for successful embryonic implantation and subsequent maintaining of pregnancy ^3^. Fundamentally, endometrium is structurally comprised of two distinct layers, outer functional layer and underlying basal layer proximal to the myometrium. It is dynamic and cyclic significant cell turnover during reproductive age ^4^. However, once normal endometrium is damaged by invasive interventions as curettage, hysteroscopic surgery or uterine artery embolization for gynecological diseases several times, endometrium becomes thin and fibrotic, leading to the disturbance of normal conception, called Asherman’s syndrome. Although several treatments for this disease, including hysteroscopic endometrial adhesiolysis and intrauterine injection of granulocyte-colony stimulating factor or platelet-rich plasma have been in use, their therapeutic efficacy is quite limited and relapse often occur after the treatment ^5,6^. Therefore, a novel therapeutic approach needs to be established for repair of endometrial receptivity when endometrium is injured.

According to recent studies, stem cell therapy using mesenchymal stem cells (MSCs) may be the breakthrough needed for the treatment of Asherman’s syndrome ^6-8^. MSCs are heterogeneous subsets of stromal/mesenchymal regenerative cells, usually derived from bone marrow, adipose tissue, dental pulp, umbilical cord and amniotic membrane ^9^. Since they possess a powerful self-renewal potency and a high ability of immunomodulation, wound healing and neovascularization, they have been widely studied as a novel stem cell resource for treatments to various diseases related with immune rejection, autoimmunity, degeneration and severe inflammation ^10-12^. Due to their regenerative ability, previous pre-clinical studies reported that intrauterine transplantation of MSCs from bone marrow, adipose tissue and umbilical cord ameliorate thin endometrium and improve fertility ^8,13-15^. However, since these MSC resources have concerns such as invasiveness of collection, low availability, minimal cost-effectiveness and ethical issues,^5,7^ novel MSC resources are warranted.

In recent years, it has been shown that endometrium contains various types of stem cells including MSCs, and these cells play an important role in dynamic endometrial regeneration during the regular menstruation cycle ^4,16,17^. Endometrial MSCs are located in the perivascular region in the functional layer. Hence, numerous endometrial MSCs shed from the endometrium into menstrual blood during menstruation, and are called menstrual blood-derived stem cells (MenSCs), first reported in 2007 ^18,19^. Therefore, menstrual blood is attractive as a novel noninvasive, periodic and cost-effective MSC resource with few ethical concerns ^20^. Until now, several pre-clinical studies showed the regenerative efficacy of MenSCs on infertility caused by injured endometrium;^21-23^ however, these previous reports have some limitations. First, because they used MenSCs only from healthy volunteers,^21-25^ the efficacy of collecting and using MenSCs from infertile patients is questionable. Second, the protocol for establishment of an injured endometrium animal model is controversial and employ ethanol intrauterine injection^13,14,26^ and heat radiation^27^ and are thought to be far from the natural etiology of thin endometrium. Third, few reports have studied the safety concerns of intrauterine transplantation of MenSCs, toxicity and tumorigenicity ^28-30^. To overcome these limitations and provide proof of concept, further studies are needed to validate whether MenSCs from infertile patients may be used as a safe stem cell resource using a clinically relevant model with injured endometrium.

In this study, we successfully collected MenSCs from infertile patients, even from a patient with Asherman’s syndrome and established a clinically relevant model with mechanically injured endometrium. Further, we show that intrauterine transplantation of MenSCs can ameliorate injured endometrium through their paracrine effects on angiogenesis and tissue repair without apparent safety concerns. Consequently, we conclude that MenSCs from infertile patients may be a promising stem cell resource, and intrauterine transplantation of autologous MenSCs may be a novel therapeutic option for endometrial infertility in the future.

## Methods

### Isolation and culture of menstrual blood-derived cells

Menstrual blood-derived cells were obtained from infertile patients recruited at our hospital with written informed consents (approval number: 2021-025). When the patients received a pelvic examination during a menstrual cycle, the clinicians had collected menstrual blood samples into a collection tube using a syringe or cotton with Dulbecco’s modified Eagle’s medium (DMEM)-high glucose (Gibco, USA, catalog number 11965-118) containing 2% fetal bovine serum (FBS) (Gibco, USA, catalog number 16000-044) and 1% penicillin-streptomycin amphotericin B suspension (AA) (FUJIFILM, Japan, catalog number 161-23181) and stored at 4°C (Supplemental Figure 1). The collection method was selected by clinicians depending on the amount of menstrual blood in patients’ vagina. After centrifugation at 1,000 rpm for 5 min, the supernatant was removed and additional 1 ml of DMEM/2% FBS/1% AA was added into the pellet. Then, 10 ml of 1X RBC lysis buffer (pluriSelect, GER, catalog number 60-00050-12) was added and gently pipetted several times. Then, the sample was incubated at 4°C for 12 min. After incubating, the mixture was gently pipetted several times and centrifuged at 1000 rpm for 5 min. Then, the supernatant was removed and rinsed with 5 ml of phosphate-buffered saline (PBS) and centrifuged at 1000 rpm for 5 min two or three times until the mixture looked transparent. The pellet was dispensed with DMEM/10% FBS/1% AA and seeded on Bio-coated culture dishes (Corning, USA, catalog number 354450) and incubated at 37°C, 95% air and 5% CO_2_. The medium was replaced at day 1 to remove non-adherent cells and afterwards every 2-3 day with DMEM/10% FBS/1% penicillin streptomycin (PS) (Gibco, USA, catalog number 15140-122). These cells were passaged serially when cells reached to 80-100 % confluency or when proliferation essentially ceased by using 0.25% Trypsin/EDTA (Gibco, USA, catalog number 25200-056) and frozen with STEM CELLBANKER (NIPPON ZENYAKU KOGYO CO., LTD., Japan, catalog number CB045) in -80°C.

We used menstrual blood-derived cells from volunteers as a control with written informed consent (approval number: 89). Menstrual samples on volunteers’ sanitary pad were collected into a collection tube or bottle with DMEM/2% FBS/1% PS by themselves and stored at 4°C. Then, the following procedure was the same as that in infertile patients.

### Tri-lineage differentiation of menstrual blood-derived cells

To evaluate the potential of MSCs, we evaluated menstrual blood-derived cells according to the criteria of the international society for cellular therapy (ISCT) ^31^. Tri-linage differentiation of cultured cells into adipocytes, osteocytes and chondrocytes was evaluated using a human mesenchymal stem cell functional identification kit (R &D systems, USA, catalog #SC006) according to the manufacturer’s instructions. Adipogenesis, osteogenesis and chondrogenesis were confirmed by immunochemistry for fatty acid binding protein-4 (FABP-4), osteocalcin and aggrecan, respectively.

### Immunochemistry

For immunocytochemistry, culture cells were fixed with 4% paraformaldehyde (PFA) in PBS for 10 min at 4°C. After washing with PBS, they were treated with 0.1% Triton X-100 (Sigma-Aldrich, #T8787-100 ML) or PBS for 10 min at 4°C, depending on the targeted antigen localization. After washing with PBS, cells or tissue sections were incubated with Protein Block Serum-Free Ready-To-Use (Dako, #X 0909) for 30 min at room temperature, followed by reaction with primary antibody in blocking buffer over night at 4°C. After washing with PBS, the sections were incubated with fluorescent-conjugated secondary antibody. Anti-rabbit or anti-goat immunoglobulin G (IgG) bound to Alexa 488 or 546 was incubated in blocking buffer for 30 min at room temperature. The nuclei were stained with -Cellstain®-DAPI solution (1:1000, DOJINDO, Japan, D523). For immunohistochemistry, the tissue sections embedded into paraffin were deparaffinized and washed with PBS. Slides were incubated with primary antibodies, followed by further incubation with horseradish peroxidase (HRP)-conjugated goat anti-mouse IgG (ZSJQ-Bio, Beijing, China, 1:100). The antibody stains were developed by the addition of diaminobenzidine (DAB) and the nucleus was stained with hematoxylin. All images were captured using fluorescence microscopy (BZ-X700, KEYENCE). Antibody information is provided in Supplemental Table 1.

### Flowcytometry

For the analysis of phenotypic markers as MSCs, cells at passage 4-7 were trypsinized and 0.3-1.0 × 10^6^ cells were dispensed into 1.5-ml tube, washed with PBS containing 3% FBS. After centrifugation at 1000 rpm for 5 min, Zombie Violet™ Fixable Viability Kit (1:100, BioLegend, catalog number 423114) was used to remove dead cells for 20 min at room temperature according to the manufacturer’s protocol. After centrifugation at 1000 rpm for 5min, cells were incubated with the specific labelled antibodies for CD 14, CD19, CD34, CD45, CD73, CD90, CD105 and human leukocyte antigen (HLA)-DR for an hour on ice (Supplemental Table 1). In all experiments, isotype antibodies were also used as negative controls. Almost 20,000 events were collected on Cell Sorter (SONY, Japan, SH800) and analyzed using Flowjo 10.8.1 software.

### Cell growth curves

Cells at passage 4-7 were plated into 6 well plates (1 × 10^5^/well). The medium was replaced every 2 or 3 days. At day 1, 3, 5 and 7 after the plating, phase-contrast photo micrographs were taken and total numbers of cells were counted by Vi-CELL XR Cell Viability Analyzer System (BECKMAN COULTER, USA,catalog number 383556).

### Collection of conditioned media and proteomics analysis

To collect conditioned media (CM) from menstrual blood-derived cells from an infertile patient and a volunteer, we followed a previous report.^32^ Briefly, these cells were seeded at 3,000 cells/cm^2^ on T25 flasks and propagated until 80% confluent. These cells were incubated for 24 h with DMEM/2% FBS. The medium was changed to M199 medium (Sigma–Aldrich) and 0.5% bovine serum albmin (BSA) (Sigma–Aldrich) and these cells were also incubated for 48 h at 37°C, 95% air and 5% CO_2_. Then, CM were collected, centrifuged at 1000 rpm for 5 min, and stored in aliquots at -80°C.

Then, to evaluate proteome of the CM, liquid chromatography tandem mass spectrometry (LS-MS) analysis was conducted at APRO Science Institute (Tokushima, Japan). Briefly, prior to the analysis, affinity-based enrichment using the agarose-immobilized benzamidine was performed to remove high abundant proteins mainly derived from basal medium ^33^. After the digestion of bound proteins from affinity beads, the quantification of peptide was performed using Pierce Quantitative Colorimetric Peptide Assay (Thermo Fisher Scientific). Then, samples were precipitated using trichloroacetic acid, followed by reduction, alkylation with iodoacetamide, and trypsinization using an equal amount of peptide between samples. The digested samples were prepared for LS-MS analysis by GL-Tip SDB (GL Science). Further, in each sample, 500 ng of peptide were analysed using Q Exactive Plus (Thermo Fisher Scientific) coupled on-line with a capillary high-performance liquid chromatography (HPLC) system (EASY-nLC 1200, Thermo Fisher Scientific) to acquire MS/MS spectra. Data derived from the MS/MS spectra were used to search the protein database SWISS-Prot using the MASCOT Server [http://www.matrixscience.com] and to identify proteins using the program Scaffold viewer [http://www.proteomesoftware.com/products/scaffold].

### Scratch assay

The effect of CM on migration of endometrial stromal cells was assessed by a scratch assay ^23,27^. For this assay, we used endometrial stromal cells at passage 6-8 derived from specimens without any abnormalities or malignancies obtained from women of reproductive age undergoing hysterectomy for benign gynecological diseases after written informed consent in a previous study approved by the Institutional Review Board of the National Center for Child Health and Development of Japan (approval number: 2289) and The Jikei University School of Medicine (approval number: 28-083(8326)) ^34^. At 100% confluence in a 96 well plate, the monolayer was scratched with Essen®□ 96-well WoundMaker™ (Essen Bioscience, inc., USA), washed with M199/0.5% BSA 3 times and then incubated under the following conditions: 10-fold diluted CM from an infertile patient and a volunteer and M199/0.5% BSA as a negative control for 24 h. Images were obtained every hour. Wound confluence (%) and relative wound density (%) was calculated by IncuCyte™ software (Essen Bioscience, inc., USA) according to the manufacturer’s instructions. Briefly, wound confluence measurement represents the fractional area of the wound that is occupied by migrated cells and relative wound density is a measurement of the density of the wound region relative to the density of the cell region. Each experiment was performed in triplicate.

### Tube forming assay

The procedure was followed and modified as previously reported ^35^. Briefly, human umbilical vein endothelial cells (HUVECs, ATCC, Cat# CRL-1730) were incubated with endothelial cell medium (ECM) (Sciencell Research Laboratory, CA, 1001) at passage 6-8 when the cells reached to 80-100% confluency. Then, the cells were trypsinized and resuspended in the following conditions: 10-fold diluted CM from an infertile patient and a volunteer, M199/0.5% BSA as a negative control and M199/0.5% BSA with recombinant VEGF (5 ng/ml) as a positive control. The cells were then seeded in 96 well plates (Corning, USA) pre-coated with 30 μl growth factor reduce (GFR) Matrigel (Corning, USA, 354230) at a density of 2 × 10^4^ cells/well. After 18-20 h, gels were stained with Calcein AM (Corning, USA,) to quantify covered area using fluorescence microscopy (BZ-X700, KEYENCE, Japan). Each experiment was performed in triplicate.

### Histological analysis

The animal samples were collected and examined histologically. For hematoxylin and eosin (H&E) stain, sections were stained with Carrazi’s hematoxylin solution (Muto Pure Chemicals Co., Ltd., Tokyo, Japan) for 5 min and 1% eosin solution (Muto Pure Chemicals Co., Ltd., Tokyo, Japan) for 30 s. To evaluate fibrosis, we used masson trichrome (MT) stain. Briefly, after stained with Weigert’s iron hematoxylin (Muto Pure Chemicals Co., Ltd., Tokyo, Japan) for 5 min, sections were stained with following solutions; 0.75% Orange G solution for 1 min (Muto Pure Chemicals Co., Ltd., Tokyo, Japan), Masson B solution (Muto Pure Chemicals Co., Ltd., Tokyo, Japan) for 20 min and then aniline blue (Muto Pure Chemicals Co., Ltd., Tokyo, Japan) for 6 min. For quantitative trichrome analysis, images of whole horizontal uterine section were electronically scanned at maginification of 200×, subsequently analyzed using Image J software. Additionally, for quantitative immunochemistry analysis, three random sections of trichromatic slides were also electronically scanned at magnification of 400× analyzed using Image J (version 1.32j) software (National Institutes of Health, USA http://rsb.info.nih.gov/ij/).

### Animal experiments

Seven-week-old female Crlj:CD1-Foxn1^nu^ ICR nude mice were purchased (The Jackson Laboratories Japan, Inc. Japan). Five-week-old male Crlj:CD1-Foxn1^nu^ ICR nude mice were also purchased for fertility testing. All mice were used for each experiment after at least a week acclimation. Mice were housed one to four mice per cage at room temperature in a specific pathogen-free condition on a 12-hour light/dark cycle and with free access to food and water.

#### Establishment of a mechanically endometrial injured model

Surgical procedures were performed in the proestrus or estrus cycles determined by vaginal smears. Mice were sterilized and anesthetized by 2-3% isoflurane inhalation, and an abdominal midline incision was made to expose both uterine horns. Then, a 20-gauge needle was inserted from a cephalic side of a uterine horn and endometrial curettage was performed at both uterine horns until the uterus became pale to the naked eye. The sham group underwent an abdominal midline incision alone without endometrial curettage. At day 0, 1, 4 and 7, the mice were euthanized and the uterine horn was harvested for the histological analysis (n=3, at each different time point in both groups). The endometrium was histologically evaluated in horizontal section. Endometrial thickness was measured by the maximal distance from the endometrial–myometrial interface to the endometrial surface (H&E stain). Endometrial area was calculated as the total area of the endometrium (H&E stain). All parameters were measured using Image J software.

#### Intrauterine transplantation of menstrual blood-derived cells

Immediately after endometrial curettage, menstrual blood-derived cells (1.0 × 10^6^ cells in 30 μl of PBS) were injected into both uterine horns of the endometrial injured model. At day 1, 7 and 30, the mice were euthanized and the uterine horns were harvested for histological analysis to evaluate intrauterine distribution of transplanted menstrual blood-derived cells. The endometrium was evaluated in the sagittal sections (n=3, at each different time point). Further, to evaluate systemic biodistribution and tumorigenicity of menstrual blood-derived cells,^36^ other major organs including brain, heart, lung, liver, spleen, pancreas, kidney, adrenal gland and ovary were also histologically evaluated.

To further evaluate the therapeutic effect of intrauterine transplanted menstrual blood-derived cells on the injured endometrium, mice were divided into 3 groups as follows: sham group, injured group and MenSC group (n=4 in each group). All groups were established as previously described above. Additionally, the injured group received 30 μl of PBS after the curettage. At day 7, mice were euthanized and the left uterine horns were collected into 20% formalin for the histological analysis, whereas the right uterine horns were collected into RNAlater™ Solution (invitrogen, USA, AM7021) for real-time quantitative polymerase chain reaction (RT-qPCR). In addition, to examine the toxicological analysis of menstrual blood-derived cells transplantation. All mice were weighed prior to the surgery at day 0 and after sacrifice at day 7, and 1 ml of blood was sampled from the inferior vena cava for blood cell counts and biochemical examination. Further, the other major organs were also weighed on day 7.

### Fertility testing

After a week from the surgical procedure, the female mice were bred with the sexually mature male mice at 1:1 ratio for a week. The morning of vaginal plug presence was recorded as gestational day 0.5 (E 0.5). After delivery, the number of litters and gestational periods were observed in the sham (n=5), injured (n=5) and MenSC groups (n=10).

### Real-time quantitative polymerase chain reaction

This procedure was followed and modified as previously reported.^34^ Briefly, RNA was extracted from homogenized tissues using ISOGEN (NIPPON GENE, Japan, 311-02501). An aliquot of total RNA was reverse-transcribed using qScript™ cDNA SuperMix (Quantabio, USA, 95048-025). For the thermal cycle reactions, the cDNA template was amplified (Applied Biosystems Quantstudio 12 K Flex Real-Time PCR System) with gene-specific primer sets (Supplemental Table 2) using the Platinum SYBR Green qPCR SuperMix-UDG with ROX (Invitrogen, #11733-046) under the following reaction conditions: 40 cycles of PCR (95°C for 15 s and 60°C for 1 min) after an initial denaturation (95°C for 2 min). Fluorescence was monitored during every PCR cycle at the annealing step. In this analysis, we targeted mRNA expression of *vascular endothelial growth factor (VEGF), fibroblast growth factor-1 (FGF-1), FGF-2* and *epidermal growth factor (EGF)* and mRNA expression levels were normalized using *glyceraldehyde-3-phosphate dehydrogenase (GAPDH)* as a housekeeping gene. Each experiment was performed in triplicate.

### Statistical analysis

Continuous data were presented as mean ± standard error of the mean (SEM) or standard deviation (SD) and analyzed using Student’s t-test. Categorical data were presented as percentages and analyzed using Chi-square test. Prism 9.4.0 software (GraphPad, Inc.) was used for statistical analyses. P < 0.05 was considered statistically significant for all analyses.

## Results

### Successful identification of mesenchymal stem cells from menstrual blood

We gathered total 5 menstrual samples from volunteers and were able to isolate menstrual blood-derived cells in all samples. In addition, we collected menstrual blood samples from a total of 16 infertile patients and successfully isolated menstrual blood-derived cells from 10 samples, even from Asherman’s syndrome patients (Table 1 and Supplemental Table 3). We failed to isolate menstrual blood-derived cells from the other 6 samples, which were mainly collected by cotton and consisted of cells that resembled vaginal epithelial cells when seeded on dishes. Following cell culture, spindle-shaped fibroblast-like cells were found from both the volunteers and the infertile patients (Fig 1A). Then, we randomly selected 3 samples from volunteers and infertile patients to determine the cellular characteristics as MenSCs. In tri-lineage differentiation, menstrual blood-derived cells from both volunteers and infertile patients expressed FABP-4 for adipogenesis, osteocalcin for osteogenesis and aggrecan for chondrogenesis (Fig 1B). Additionally, upon flowcytometric analysis for evaluating phenotypic markers of MSCs, these samples expressed CD73, CD90 and CD105 as MSC markers without the expression of CD14, CD19, CD34, CD45 and HLA-DR as hematopoietic cell markers (Fig 1C). Expression of these MSC markers was similar between menstrual blood-derived cells from volunteers and infertile patients (Supplemental Table 4). Further, we also confirmed the cells with characteristics of MenSCs from an Asherman’s syndrome (patient #1) (data not shown). These results show that we could identify menstrual blood-derived cells fulfilling the ISCT criteria from volunteers and infertile patients.

**Table 1.** Patient characteristics of samples with successful primary culture of menstrual-blood derived cells

| Sample # | Age, y | Gravida | Parity | BMI, kg/m<sup>2</sup> | Cause for infertility | Duration of infertility, y | Treatment method | Menstrual day | Collection method |
| --- | --- | --- | --- | --- | --- | --- | --- | --- | --- |
| 1 | 40 | 3 | 1 | 19.3 | Asherman's syndrome | 1.3 | TI | D2 | Syringe |
| 2 | 39 | 2 | 1 | 23.4 | Ovulation disorder | 0.5 | IUI | D3 | Syringe |
| 3 | 39 | 1 | 1 | 19.9 | Ovarian insufficiency | 1.5 | ART | D2 | Cotton |
| 4 | 40 | 2 | 0 | 20.9 | Endometriosis | 0.7 | ART | D2 | Syringe |
| 6 | 28 | 0 | 0 | 23.8 | Unexplained | 0.5 | IUI | D2 | Syringe |
| 8 | 38 | 1 | 1 | 19.5 | Hydrosalpinx, Luteal insufficiency | 5.0 | ART | D3 | Cotton |
| 9 | 41 | 1 | 1 | 24.7 | Unexplained | 0.4 | IUI | D2 | Syringe |
| 10 | 45 | 3 | 1 | 22.4 | Unexplained | 0.1 | ART | D3 | Cotton |
| 13 | 38 | 1 | 0 | 25.1 | Unexplained | 3.3 | ART | D4 | Cotton |
| 14 | 46 | 1 | 1 | 17.7 | Ovarian insufficiency | 7.0 | TI | D2 | Syringe |

y, year; BMI, body mass index; TI, timed intercourse; IUI, intrauterine insemination; ART, assisted reproductive technology; D, day.

**Figure 1.**
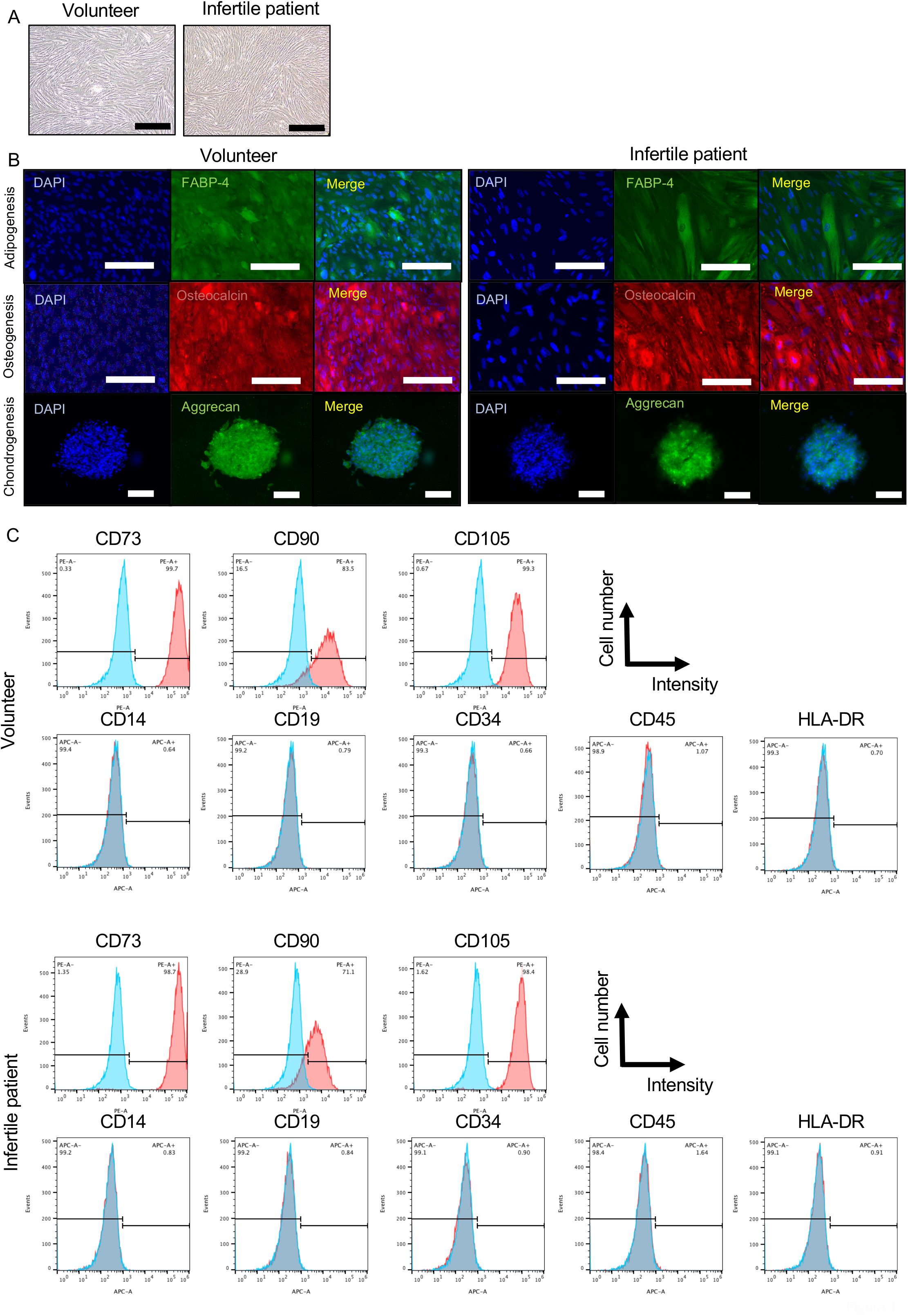
Cellular characterization of menstrual blood-derived cells as mesenchymal stem cells. (A) Phase-contrast photomicrographs of spindle-shaped fibroblast-like menstrual blood-derived cells from volunteers and infertile patients with 100% confluency on a petri dish. Black bar is 500 μm. (B) Immunocytochemistry of menstrual blood-derived cells after differentiation. Menstrual blood-derived cells from both volunteers and infertile patients were positive for FABP-4 (green), osteocalcin (red), and aggrecan (green) for adipogenesis, osteogenesis, and chondrogenesis, respectively. Cell nuclei were stained using DAPI in blue. White bar is 100 μm. (C) Flowcytometric analysis for phenotypic markers of menstrual blood-derived cells. Menstrual blood-derived cells at passage 3-4 from both volunteers and infertile patients were positive for CD73, 90 and 105 as makers for mesenchymal stem cells, and negative for CD14, CD19, CD34, CD45 and HLA-DR as makers for hematopoietic stem cells (n=3 in each group).

MenSCs from infertile patients were biologically equivalent to those from volunteers To investigate therapeutic potential equivalence between volunteer-derived and infertile patient-derived cells, we first examined the proliferative potential of MenSCs. MenSCs from both volunteers and infertile patients were propagated to 100% confluency until day 7 after plating on a 6 well plate (Fig 2A). Growth of primary menstrual blood-derived cells from infertile patients was similar to that of volunteers (Fig 2B). Further, to investigate the effect of the freeze-thaw procedure on cell growth, we compared cell proliferation between primary and thawed frozen menstrual blood-derived cells from infertile patients (Supplemental Figure 2A, B). The thawed cells were also propagated to 100% confluency at day 7 and there were no significant differences until day 7, compared to the primary cells, suggesting that proliferative potential is preserved even after the freeze-thaw procedure.

**Figure 2.**
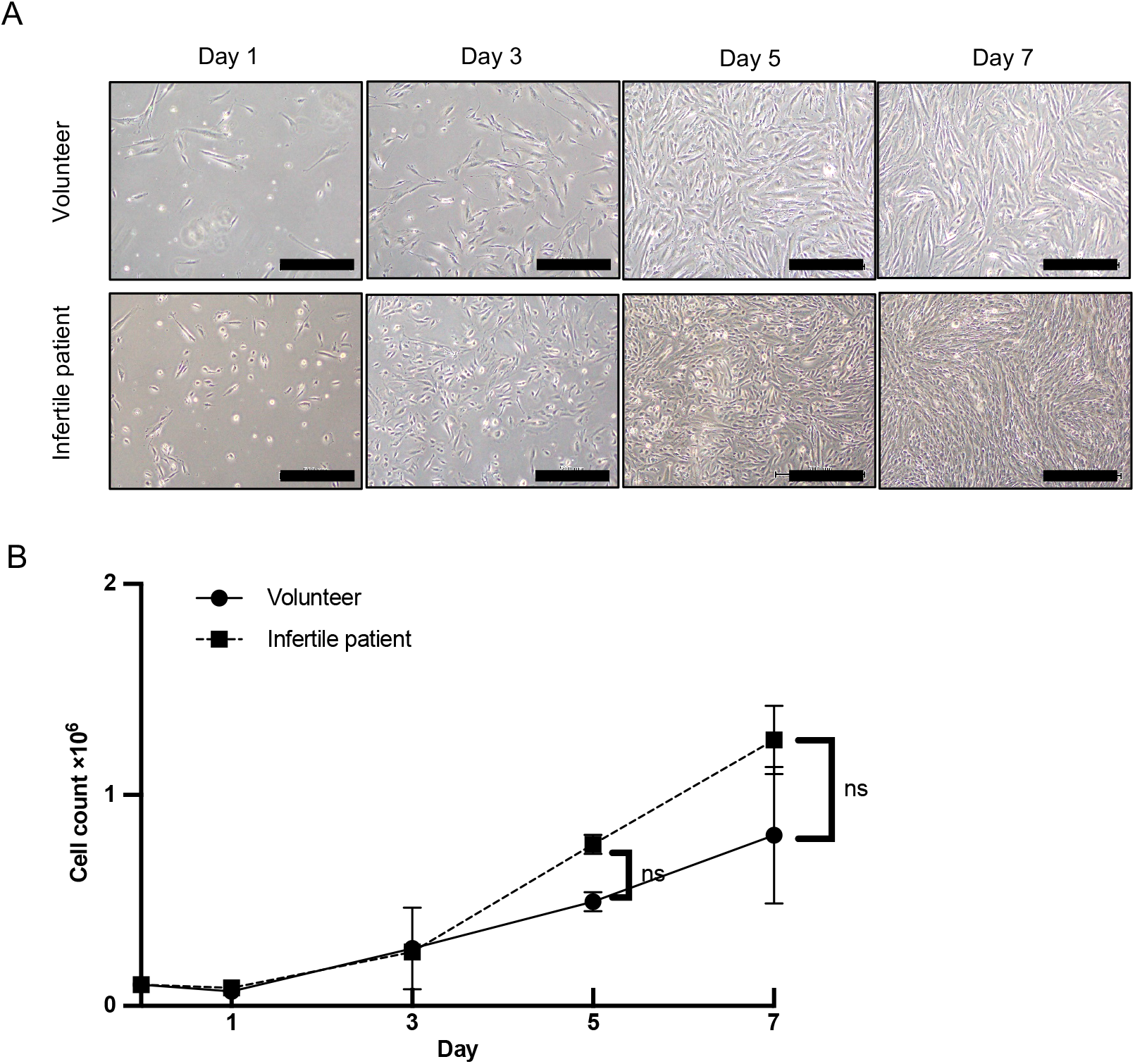
Proliferative capacity of MenSCs from volunteers and infertile patients. (A) Phase-contrast photomicrographs of MenSCs at different time points. (B) Primary cells (1.0 × 10^5^) from volunteers and infertile patients at passage 4-7 were plated into 6-well plates (n=3, in each group). Proliferative capacity was not significantly different between MenSCs from volunteers and infertile patients. Statistical significance is shown as *P <0.05 and **P <0.01. ‘ns’ means ‘not significant’. Black bars are 500 μm.

To screen comprehensively for paracrine factors from menstrual blood-derived cells, CM were collected and assessed by proteomic analysis (n=1). This analysis revealed more than 200 proteins in the CM. Quantitative scatterplot indicated that the amounts of proteins were similar between the volunteer-derived and infertile patient-derived CM (Supplemental Fig 3A). Furthermore, the number of ‘unique proteins’ (Supplemental Fig 3B) was similar in the volunteer-derived and infertile patient-derived CM in the categories of ‘biological process’, ‘cellular component’ and ‘molecular function’. In a Venn diagram, 51 proteins common in CM (Supplemental Fig 3C) were categorized into subgroups of ‘biological process’ such as ‘cellular process’, ‘biological regulation’, ‘developmental process’, ‘multicellular organismal process’, ‘response to stimulus’, ‘localization’, and ‘biological adhesion’ (Supplemental Fig 3D). In GO term analysis, 9 angiogenesis-related proteins, i.e. thrombospondin 1, fibronectin, lysyl oxidase homolog 2, junction plakoglobin, myosin-9, sushi repeat-containing protein, annexin A2, 14-3-3 protein zeta/delta and heat shock protein beta-1 were found in both groups; Cell proliferation-related 5 proteins, i.e. thrombospondin 1, fibronectin, actin beta, endosialin and c-type lectin domain family 11 member A, were common; fibroblast proliferation-regulated proteins, i.e. fibronectin, annexin A2 and endosialin, were common; cell adhesion-related 13 proteins, i.e. thrombospondin 1/2, fibronectin, collagen alpha-1(XII) chain, nidogen-1/2, lysyl oxidase homolog 2, junction plakoglobin, amyloid-beta precursor protein, laminin subunit gamma-1/beta-1, galectin-3-binding protein, testican-1, versican core protein and filaggrin-2 were in both groups. Proteomic analysis showed that infertile patient-derived MenSCs had paracrine factors involved in angiogenesis and wound healing in common with volunteer-derived MenSCs.

We then performed a scratch assay to investigate regenerative ability in injured tissue. The area of the scratch wound in each group was gradually masked by stromal cell migration (Fig 3A). Wound confluency of infertile patient-derived and volunteer-derived CM groups increased, compared to a negative control (Fig 3B). Additionally, relative wound density of infertile patient-derived and volunteer-derived CM groups also significantly increased, compared to a negative control (Fig 3C). To investigate angiogenic potential, we performed a tube-forming assay to investigate whether HUVECs form tubular networks in the presence of the volunteer-derived and infertile patient-derived CM. The capacity of HUVECs to form tubular networks in the CM were significantly increased compared to that in the negative control and equivalent to that in recombinant VEGF-supplemented culture medium, as indicated by tube area measurements (Figure 3D, E). Further, the capacity of HUVECs to form tubular networks in the infertile-derived CM was equal to the volunteer-derived CM (Figure 3D, E).

**Figure 3.**
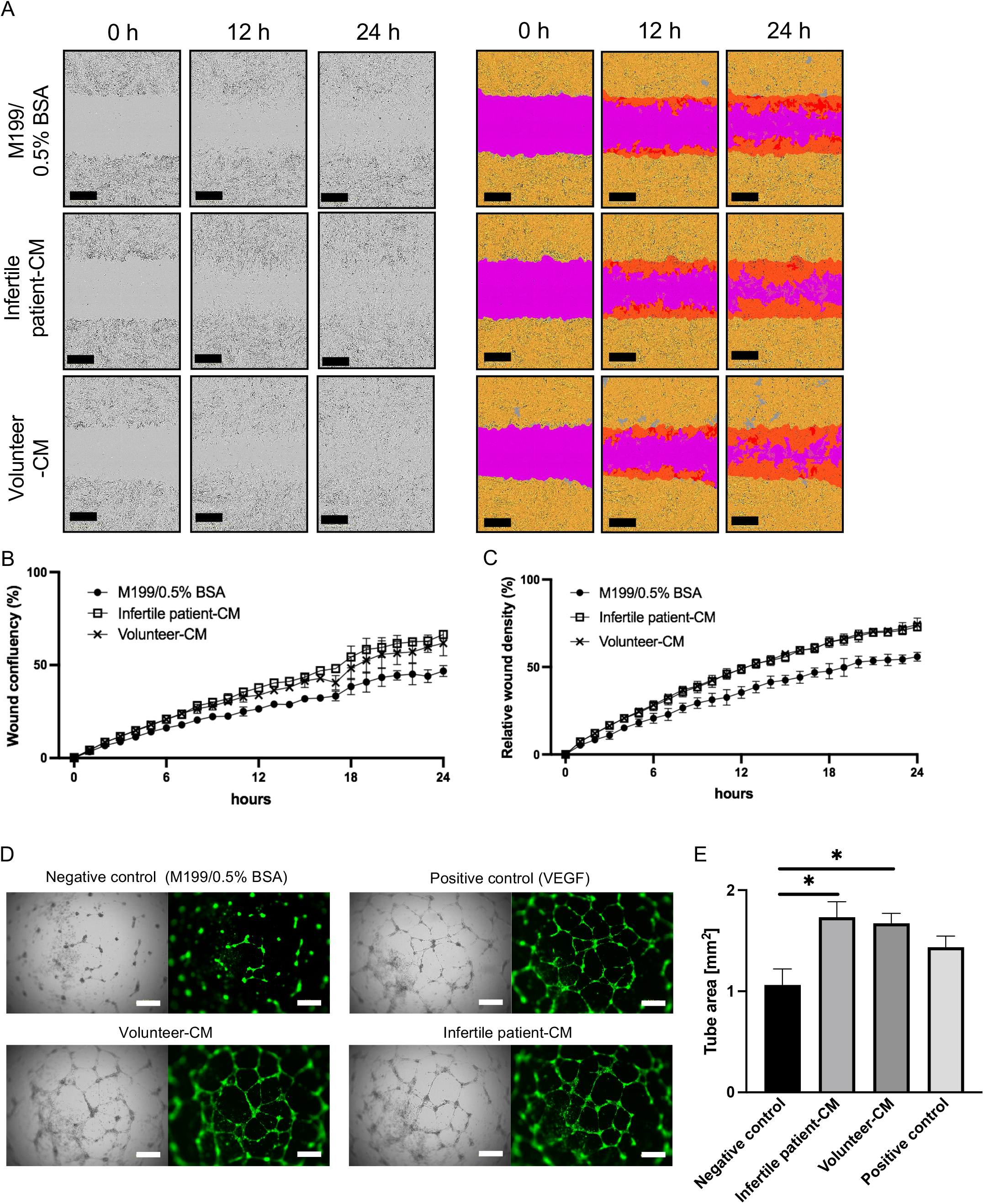
MenSCs show paracrine capacity for tissue repair and angiogenesis. (A-C) Scratch assay to evaluate the paracrine capacity for wound healing. (A) The left panels show phase-contrast photomicrographs of scratch wounds at 0, 12 and 24 h after scratching in the following media; M199/0.5% BSA (negative control), infertile patient-CM and volunteer-CM. In the right panels, the cell region is yellow, the wound region is pink and the wound mask region is orange. Black bars are 400 μm. (B) Wound confluency of infertile patient- and volunteer-derived CM groups increased, compared to the negative control (M199/0.5% BSA). (C) Relative wound density of infertile patient- and volunteer-derived CM groups significantly increased, compared to the negative control (M199/0.5% BSA). Each experiment was performed in triplicate. (D) Tube-forming assay to evaluate the paracrine capacity for angiogenesis. Microscopic appearance of tubular networks of HUVECs stained with Calcein AM in green with following media; M199/0.5% BSA (negative control), M199/0.5% BSA with recombinant VEGF (5 ng/ml) (positive control), infertile patient-CM and volunteer-CM. White bars are 500 μm. (E) The capacity of HUVECs to form tubular networks in the CM from a volunteer and an infertile patient significantly increased. Each experiment was performed in triplicate. All measurements are shown as mean ± SEM. Statistical significance is shown as *P <0.05.

These results indicate that the cellular and biological characteristic of menstrual blood-derived cells from an infertile patient were equivalent to those from a volunteer.

### Successful establishment of mechanically injured endometrium model

We performed an experiment to establish a mechanically damaged endometrium model (Fig 4A and Supplemental File 1). In comparison to the sham group, endometrial damage was apparent in the injured group at days 0 and 1 (Fig 4B, H&E). Although arrangement of endometrial epithelial and stromal cells in the injured group was repaired by day 4, their endometrial lumen looked flat (Fig 4B, H&E). Furthermore, in the injured group, fibrotic areas were found at days 1, 4 and 7 (Fig 4B, MT), while no fibrotic areas were observed in the sham group. In addition, endometrial thickness in the injured group was significantly thinner than in the sham group at day 4 and 7 (Fig 4C), whereas number of glands and endometrial area in the injured group decreased at each time point, but there were no significant differences between the sham and injured groups (Fig 4D, E). We therefore established a mechanically injured endometrium model by endometrial curettage.

**Figure 4.**
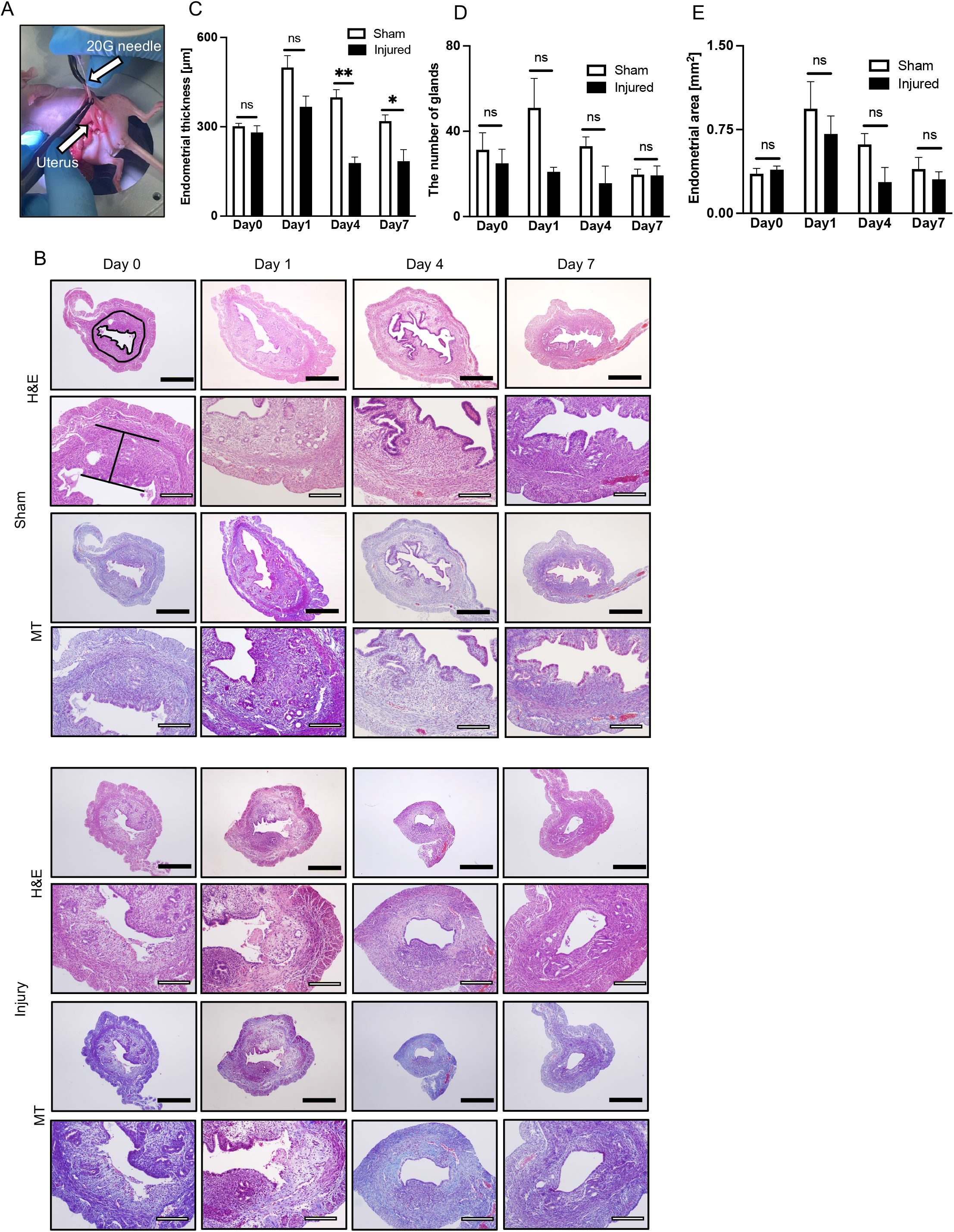
Establishment of a mouse model for mechanically injured endometrium. (A) Procedure for endometrial injury. A mouse model for endometrial injury was established by physical endometrial scratches using a 20-gauge needle in uterine horns, inserted from a cephalic side of the uterine horns. (B) Comparison of endometrial histology between sham groups and injury models at different time points after the surgical procedure (n=3 in each group). Endometrial thickness was evaluated in a sagittal section of a left uterine horn as the maximal distance from endometrial–myometrial interface to endometrial surface (shown in H&E stain of the sham group at day 0). In addition, the endometrial area was evaluated in a sagittal section of a left uterine horn as the total area of the endometrium (shown in H&E stain of the sham control at day 0). In the injured models, endometrial damage was observed by H&E staining at day 0. Further, fibrotic areas in endometrium were detected by Masson trichrome stain at day 1, 4 and 7, whereas there were no fibrotic areas observed in the sham control. (C) Endometrial thickness in the injured model was significantly thinner at days 4 and 7. (D) There was no significant difference in the number of glands at each different time point. (E) There was no significant difference in the endometrial area at each different time point. All measurements are shown as mean ± SEM. Statistical significance is shown as *P <0.05 and **P <0.01. ‘ns’ means ‘not significant’. Black bars are 500 μm and white bars are 200 μm.

### Successful establishment of an intrauterine transplantation protocol and no findings of tumorigenicity from transplanted menstrual blood-derived cells

We performed intrauterine transplantation of MenSCs to determine the intrauterine distribution of the transplanted cells (Fig 5A, B). At day 1 after transplantation, MenSCs were detected as human vimentin- and lamin AC-positive cells at the injured sites of endometrium including endometrial stroma, showing successful transplantation into injured endometrium. At day 7 after the transplantation, we found small quantity of the cells positive for human vimentin and lamin AC in endometrial stroma; however, these could not be observed at day 30. No apparent tumor formation was observed in endometrium. These findings suggest that transplanted cells gradually disappear in the mouse uterus without local tumorigenicity, probably due to engraftment failure or immunogenicity.

**Figure 5.**
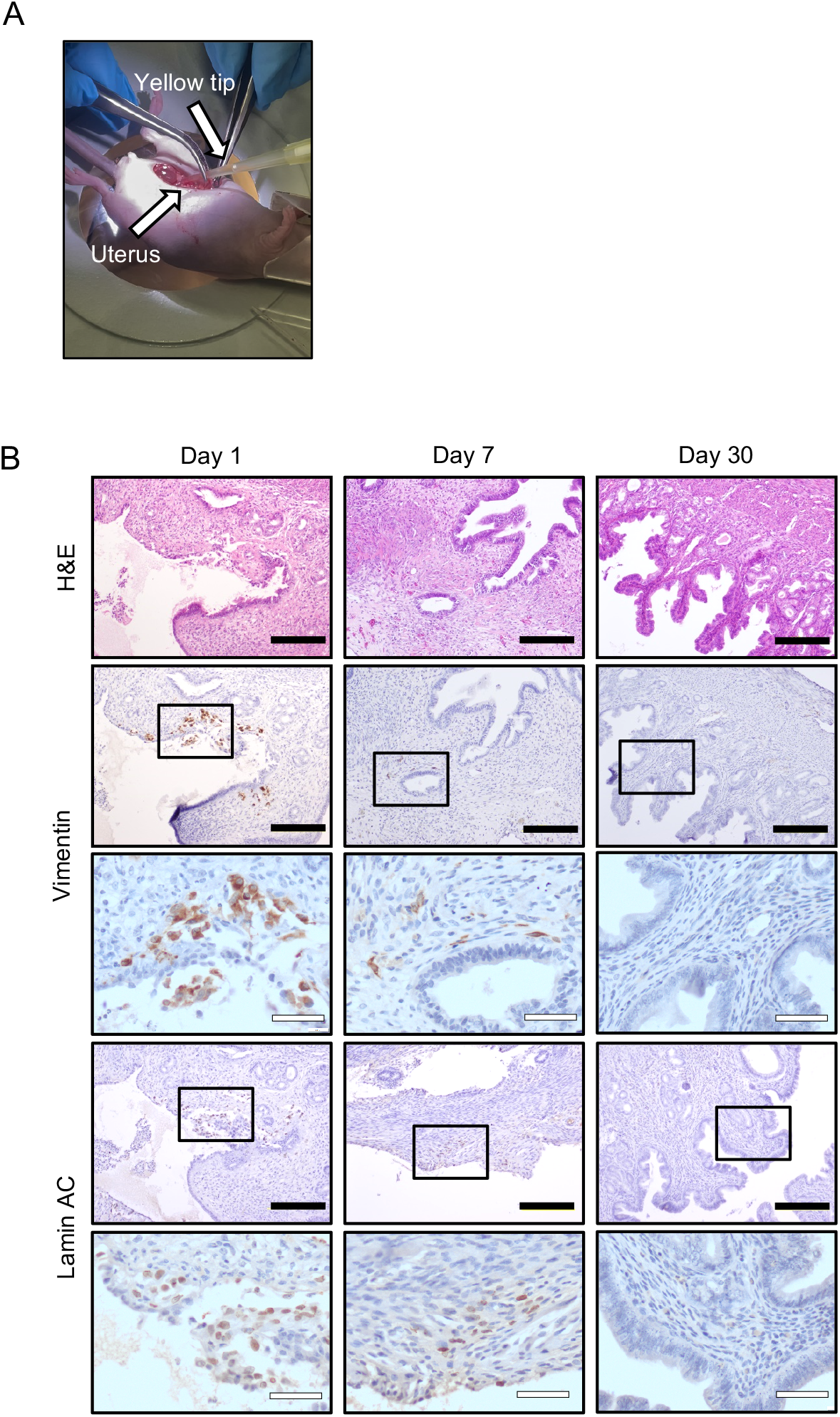
Local distribution of MenSCs after intrauterine transplant. (A) Procedure for intrauterine transplantation of MenSCs. After endometrial scratching with a 20-gauge needle, 1.0 × 10^6^ of menstrual blood-derived cells in 30 μl of PBS were injected into uterine horns. (B) Local distribution of transplanted intrauterine MenSCs at each time point (n=3). The transplanted cells were positive for human vimentin and lamin AC at day 1 in injured endometrial lesions. Small numbers of cells were found at day 7 in the stroma, not in the epithelia. At day 30, the cells positive for human vimentin and lamin AC were no longer detected. Black bars are 200 μm and white bars are 50 μm.

To assess the systemic biodistribution and tumorigenicity of the intrauterine transplanted cells, we histologically evaluated the major organs (Supplemental Figure 4). There were no obvious human vimentin-positive cells and tumor formation at 1, 7 and 30 days after intrauterine transplantation. In addition, no tumor formation was observed on the peritoneum by gross findings at the different time points.

### Menstrual blood-derived cells from an infertile patient repair injured endometrium without apparent toxicological concerns

To evaluate therapeutic effect of infertile patient-derived MenSCs on injured endometrium, we performed histopathological and RT-qPCR analysis of mice uteri at day 7 after intrauterine transplantation. The uterus in the MenSC group looked thicker, like the sham group on gross examination, whereas that in the injured group looked atrophied and congested (Fig 6A). Uterine weight in the MenSC group was slightly heavier compared to the injured group, without a statistically significant difference (Fig 6B). The endometrial thickness in the MenSC group was significantly thicker than the injured group (Fig 6C). The number of glands in the MenSC group slightly increased, compared to the injured group (Fig 6D). The endometrial area in the MenSC group was significantly larger than the injured model (Fig 6E). Further, the endometrial fibrotic area in the MenSC group was significantly decreased compared to the injured group (Fig 6F). We performed immunohistochemistry to estimate cell proliferation and neovascularization by using an antibody to Ki-67 and CD34, respectively. The Ki-67-positive cells increased in the MenSC group, compared to the injured groups, without significant statistical difference (Fig 6A, G). CD34-positive microvessels significantly increased in the MenSC group, compared to the sham and injured group (Fig 6A, H).

**Figure 6.**
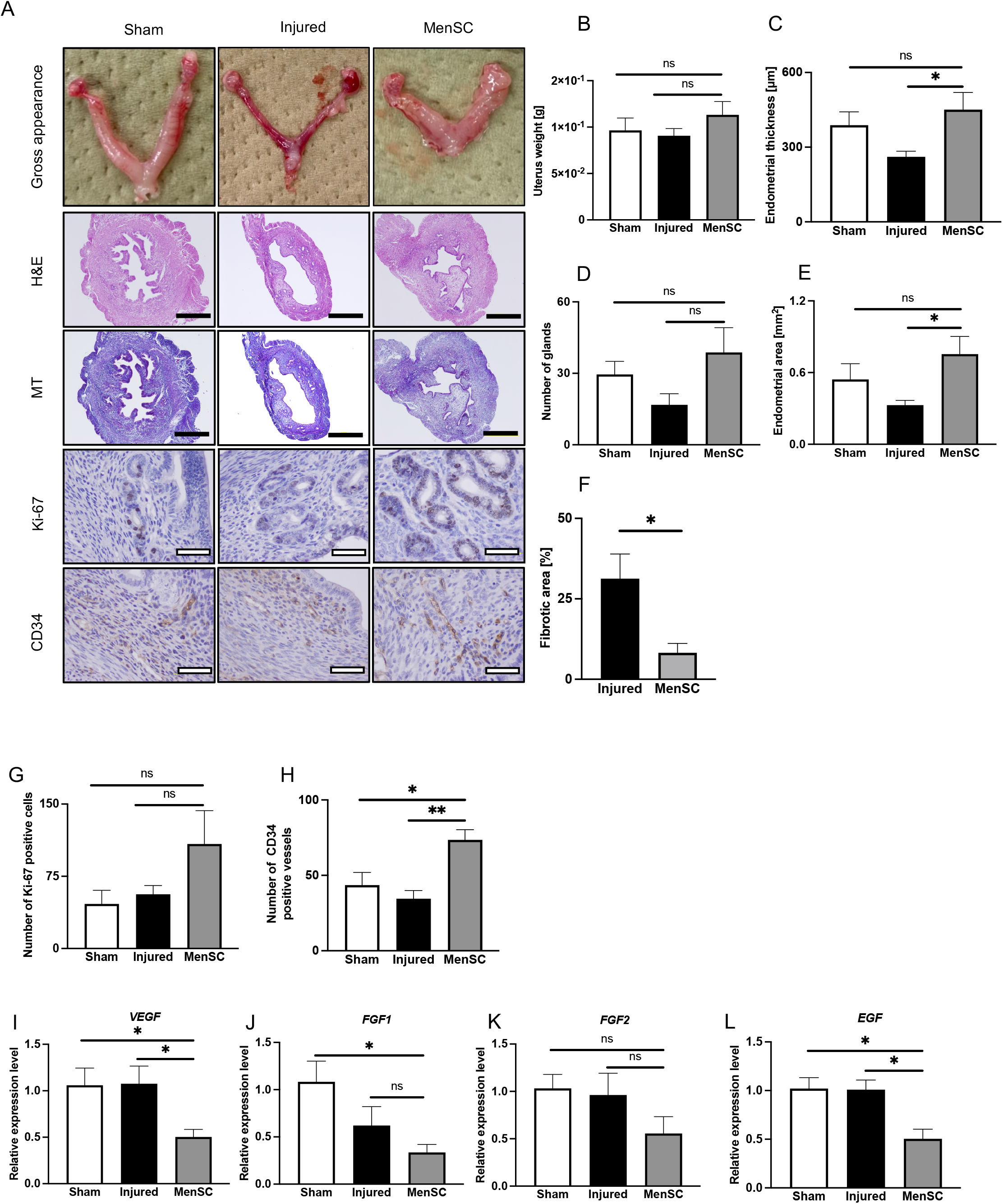
Regenerative effect of intrauterine MenSCs transplanted into injured endometrium. (A) Gross appearance and histological findings of endometrium in the sham, injured and MenSC groups at day 7 (n=4 in each group). Although the uteri of the injured group were atrophied and congested, those of the MenSC group exhibited similar gross appearance to the sham group, that is, neither atrophied nor congested. (B) There was no significant difference of uterine weights between the groups. (C) Significant recovery in endometrial thickness was observed in the MenSC group, compared to the injured group. (D) Number of glands in the MenSC group slightly increased, compared to the injured group (P = 0.10). (E) The endometrial area in the MenSC group was significantly larger than the injured group (P <0.05). (F) Endometrial fibrotic area between the injured and MenSC groups. Endometrial fibrotic area in the MenSC group significantly decreased, compared to the injured group (P <0.05). (G) Number of Ki-67 positive cells was slightly upregulated in the MenSC group, compared to the sham and injured group (P = 0.19). (H) Number of CD34-positive microvessels in the MenSC group significantly increased, compared to the injured group (P <0.01). (I-L) Gene expression of *VEGF, FGF1, FGF2* and *EGF* in each group. The expression of *VEGF, FGF1, FGF2* and *EGF* in the MenSC group significantly decreased, compared to the sham group. The expression of *VEGF* and *EGF* in the MenSC group significantly decreased, compared to the injured group (P <0.01). The experiment was performed in triplicate. Gene expression in the controls is regarded as equal to 1.0. All measurements are shown in mean ± SEM. Statistical significance is shown as *P <0.05 and **P <0.01. ‘ns’ means ‘not significant’. Black bars are 500 μm and white bars are 50 μm.

We then analyzed gene expression of whole uterine tissue in each group. The mRNA expression levels of *VEGF, FGF-1, FGF-2* and *EGF* in the MenSC group decreased, compared to the sham and injured groups. The genes for *VEGF* and *EGF* were statistically significantly decreased in the MenSC group, compared to the sham and injured groups (Fig 6I-L).

We also performed physical and blood sample examinations on each group. Between day 0 and 7 after transplantation, total body and major organ weights were not significantly different between the groups (Supplemental Table 6). Likewise, the blood cell counts and biochemical analysis showed no significant changes among the groups (Supplemental Table 7).

These results imply that intrauterine transplantation of infertile patient-derived MenSCs repairs endometrial injury and prevents fibrosis through promotion of endometrial cell proliferation and neovascularization without apparent toxicological concerns.

### Fertility testing

To further evaluate the therapeutic efficacy of the intrauterine transplantation of infertile patient-derived MenSCs for fertility, we observed the gestational period and the number of delivered litters in each group (Fig 7A). All mice in the study with vaginal plugs got pregnant and delivered pups. The gestational period was not significantly different among the sham, injured and MenSC groups (18.9 ± 0.4, 19.9 ± 0.24 and 19.3 ± 0.2 days, respectively [mean ± SEM]; P = 0.09). However, the number of pups in the injured group significantly decreased in the comparison to the sham group (7.2 ± 1.4 vs 11.6 ± 0.5 [mean ± SEM]; P <0.05). Furthermore, the average number of pups in the MenSC group increased compared to the injured group by almost 2 pups, although the difference was not statistically significant (9.1 ± 1.0 vs 7.2 ± 1.4 [mean ± SEM]; P = 0.28) (Fig 7B). These results show that the intrauterine transplantation of infertile patient-derived MenSCs partially ameliorates fertility in the mice with injured endometrium.

**Figure 7.**
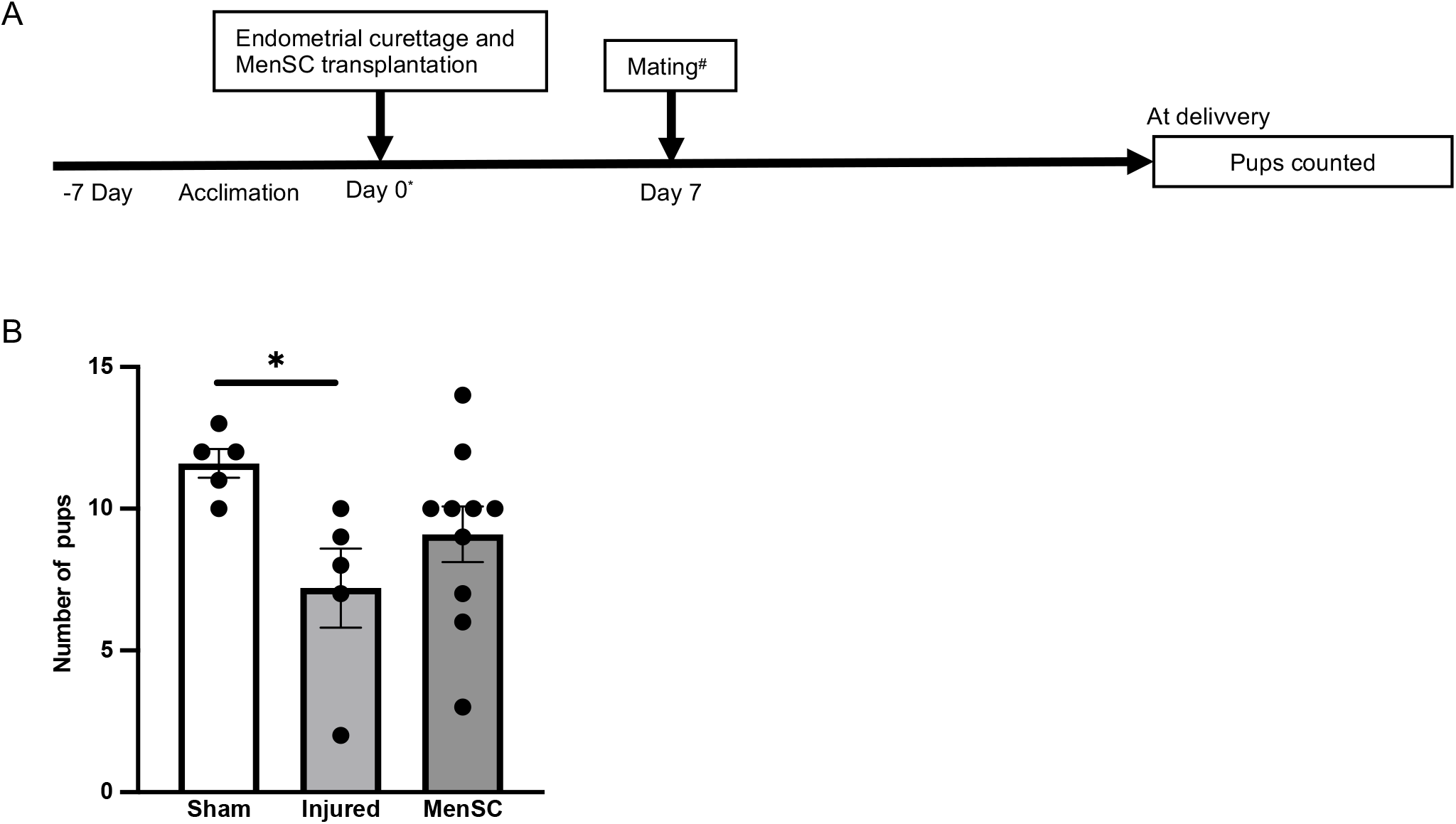
Therapeutic efficacy of transplanted intrauterine menstrual blood-derived cells for fertilization. (A) Scheme for evaluating fertility after intrauterine transplantation of MenSCs. (B) Number of pups delivered in each group. Number of pups in the injured group (n=5) significantly decreased compared to the sham group (n=5) (P < 0.05). Number of pups in the MenSCs group (n=10) partially increased compared to the injured group (P = 0.28). All measurements are shown in mean ± SEM. Statistical significance is shown as *P <0.05 and **P <0.01. ‘ns’ means ‘not significant’.

## Discussion

MenSCs have been projected to be a desirable mesenchymal stem cell resource due to their easy accessibility, periodic acquisition, beneficial cost-effectiveness, high proliferative ability, low immunological rejection and paracrine secretion of various growth and angiogenic factors ^20,28,32,37-39^. In infertility, these cells have been investigated for their potential as a novel stem-cell therapy to infertility due to thin endometrium ^21-23^. In this study, we successfully isolated and cultured menstrual blood-derived cells from infertile patients, even from Asherman’s syndrome, and demonstrated that their cellular potential as MSCs is equivalent to those from volunteers. Further, we showed their regenerative therapeutic effect on injured endometrium of the mouse model without obvious safety concerns such as toxicity and tumorigenicity, suggesting that infertile patient-derived MenSCs are a novel autologous stem cell therapy for infertility caused by injured endometrium.

### A collection method for menstrual blood-derived cells from infertile patients

Previous reports about the regenerative effect of MenSCs on injured and fibrotic organs mainly used those derived from healthy volunteers, not infertile patients ^21-25^. In such studies, menstrual cups were used to collect menstrual blood, but these cups are not widely used in Japan and hygiene issues remain. Furthermore, little is known about whether MenSCs can be collected and isolated from menstrual blood of infertile patients, particular in endometrial infertility. Therefore, other approaches should be considered to collect MenSCs. In this study, we established an efficient method for collection of MenSCs from infertile patients fulfilling with the ISCT criteria ^31^ in routine pelvic examination at infertile outpatient office, even from patients with Asherman’s syndrome. According to the protocol in this study, 10 of 16 samples were successfully cultured; on the other hand, cells could not be isolated from 6 samples. Among them, 5 of 6 samples were collected by cotton and these samples mainly contained vaginal-epithelial like cells, indicating that a syringe is preferred to cotton as a collection method. MenSCs from infertile patients may become a more acceptable stem-cell resource in the future, since our simple method to collect MenSCs can be introduced in infertility clinics.

### A clinically relevant mouse model with injured endometrium

Another problem is differential methods to establish animal models with injured endometrium in previous studies. The methods include intrauterine ethanol injection ^13,14,26^, intrauterine heat irradiation ^27^, electric coagulation ^23^, and introduction of lipopolysaccharide ^40^. Although these protocols can successfully establish thin endometrium animal models, most intrauterine adhesions occur after intrauterine mechanical intervention in the clinic ^41^; that is, these models reported in previous studies are far from the natural pathology of infertility by thin endometrium. This fact indicates that the therapeutic effect of MenSCs using these models might not be enough for acquisition of proof of concept. As other reports using a mechanically injured model indicated ^15,42,43^, the model in this study showed thinner endometrium, fibrotic lesions and impaired fertility is clinically relevant to the pathology of infertility with injured endometrium.

### Regenerative effect of menstrual blood-derived cells on injured endometrium through paracrine effects without apparent safety concerns

Recent studies show that the regenerative effect of implanted or transplanted MSCs is through a paracrine effect rather than differentiation ability ^8,44^. Conditioned media of bone marrow derived MSCs contain abundant angiogenic, immunomodulatory and growth factors for tissue repair ^45^. Additionally, MenSCs from healthy volunteers secrete high levels of angiogenic and anti-inflammatory factors ^28^. However, few reports investigated the secretary capacity of infertile patient-derived MenSCs. Our results showed that MenSCs from infertile patients secrete various factors related to angiogenesis and tissue repair, equal to volunteer-derived MenSCs. It is noteworthy that there are differences in cellular potential between the samples, though. Significant endometrial repair is probably due to neovascularization and prevention of fibrosis after intrauterine transplantation of MenSCs, without direct differentiation into endometrial epithelial or stromal cells; the regenerative potential of infertile-patient derived MenSCs is based on their paracrine effect.

Decreased expression of VEGFA, FGF-1, FGF-2, and EGF in the uteri of mice in the MenSCs group is compatible with a therapeutic effect of MenSCs on decrease of VEGFA expression and relative increase of CD34-positive vessels in a diabetic wound repair model ^46^. Adipose tissue-derived MSCs have no effect in expression of FGF-1/2, EGF and Angiopoietin-1/2 ^13^. Likewise, intravenous injection of umbilical cord-derived MSCs have no effect in expression of inflammatory factors including VEGF ^47^. Based on these reports, suppression of these cytokines occurs probably due to several reasons such as factors from MenSCs, MSC source or the experimental procedures used in each report.

We herewith emphasize that intrauterine transplantation of MenSCs has a positive therapeutical potential on injured endometrium and impaired fertility. Single-cell transcriptomatic analysis showed that the underlying pathology of Asherman’s syndrome is deficiency and senescence of endometrial perivascular cells ^48,49^ which are a main component of MenSCs ^44^. Intrauterine transplantation of autologous MenSCs is a rational approach for treating thin endometrium. Our experimental procedure, MenSC transplantation on the same day of the curettage, indicates their prophylactic efficacy on the injured endometrium. Autologous intrauterine transplantation of MenSCs may not only be a curative therapy for Asherman’s syndrome but a prophylactic therapy for patients with injured endometrium after endometrial curettage, hysteroscopic myomectomy and endometrial ablation.

Intrauterine transplantation of MenSCs appears to have no safety concerns such as toxicity and tumorigenicity ^29^. Another study reports no safety concerns on immunogenicity and tumorigenicity after intraperitoneal and intravenous transplantations of MenSCs ^28^. According to both reports, transplantation of MenSCs shows no apparent safety concerns. Since these studies only evaluate short-term complications, future studies should look for evidence of long-term complications.

### Future inspection of the regenerative therapy for injured endometrium using menstrual blood-derived cells

Several studies of phase I clinical trials using transplantation of autologous MSCs for severe Asherman’s syndrome have been reported: one for intra-artery transplantation of bone marrow-derived stem cells ^50^ and two for intrauterine transplantation of MenSCs ^51,52^. These trials showed favorable results: recovery of endometrial thickness and successful pregnancy. A clinical trial using autologous MenSC transplantation will possibly be a more effective and safer treatment for injured endometrium. However, unsolved problems still remain: what quantities of cells are most effective, how many rounds of MenSCs transplantation are required and when is the best timing for transplantation after intrauterine interventions. The effectiveness of prophylactic transplantation of MenSCs has not yet been investigated in clinical trials. In addition, some reports indicate that intrauterine transplantation of bio-engineered scaffold-embedded MSCs ^5,53^ or extracellular vesicles such as exosomes from MSCs ^6,30,54^ are preferred to MSC administration alone. Further investigation to determine the most effective therapeutic protocol using infertile patient-derived MenSCs on injured endometrium will be needed.

## Conclusion

In conclusion, our study demonstrates that autologous intrauterine transplantation of infertile patient-derived MenSCs exert their regenerative capacity as a curative and prophylactic treatment for injured endometrium without apparent safety concerns. MenSCs are an attractive and novel cell source for autologous regenerative therapy for injured endometrium. Further safety examination using good laboratory practice is needed before clinical trials.

## Declarations

### Ethics approval and consent to participate

The protocol for the use of menstrual blood-derived cells in the present study was approved by the Institutional Review Board of National Center for Child Health and Development of Japan (approval number: 2021-025 for infertile patients and 89 for volunteers). Use of endometrial stromal cells was approved by the Institutional Review Board of National Center for Child Health and Development of Japan (approval number: 2289) and The Jikei University School of Medicine (approval number: 28-083(8326)). The animal experiments were approved by the Institutional Animal Care and Use Committee of National Center for Child Health and Development (approval number: 2003-002). All animal experiments were based on the 3R principle (refine, reduce, and replace), and all efforts were made to minimize animal suffering and to reduce the number of animals used.

### Consent for publication

Not applicable.

### Availability of data and material

The datasets used and/or analyzed during the current study are available from the corresponding author on reasonable request.

## Authors’ contributions

Conceptualization: SH, RY, AU, Sample collection: EK, MA, TS, Data curation: SH, RY, YF, H Kishigami, Formal analysis: SH, RY, YF, HK, Funding acquisition: SH, RY, AU, Investigation: SH, RY, YF, H Kishigami, Methodology: SH, RY, TK, AU, Project administration: HS, AO, AU, Software: SH, RY, Supervision: H Kishi, AO, HS, AU, Writing–original draft: SH, RY, Writing–review & editing: H Kishigami, YF, EK, MA, TS, H Kishi, HS, AO, AU. All authors approved the final version of the manuscript.

## Acknowledgments

We would like to express our sincere thanks to K. Miyado and H. Akutsu for fruitful discussion, to M. Ichinose for providing expert technical assistance, to C. Ketcham for English editing and proofreading, and to E. Suzuki and K. Saito for secretarial work.

## List of Abbreviations

MSCs: Mesenchymal stem cells
MenSCs: menstrual blood-derived stem cells
DMEM: Dulbecco’s modified Eagle’s medium
FBS: fetal bovine serum
AA: penicillin-streptomycin amphotericin B suspension
PBS: phosphate-buffered saline
PS: penicillin streptomycin
ISCT: international society for cellular therapy
FABP-4: fatty acid binding protein-4
PFA: paraformaldehyde
IgG: immunoglobulin G
HRP: horseradish peroxidase
DAB: diaminobenzidine
CM: conditioned media
BSA: bovine serum albumin
LS-MS: liquid chromatography tandem mass spectrometry
HPLC: high-performance liquid chromatography
HUVECs: human umbilical vein endothelial cells
ECM: endothelial cell medium
GFR: growth factor reduce
H&E: hematoxylin and eosin
MT: masson trichrome
RT-qPCR: real-time quantitative polymerase chain reaction
VEGF: vascular endothelial growth factor
FGF: fibroblast growth factor
EGF: epidermal growth factor
GAPDH: glyceraldehyde-3-phosphate dehydrogenase
SEM: standard error of the mean
SD: standard deviation

## Supplemental figure legends

**Supplemental Figure 1. A collection method of MenSCs from infertile patients**

Collection method for menstrual blood samples. Samples were collected into 15-ml tubes with a syringe (A) or a 150-ml bottle with a cotton (B), containing DMEM/2% FBS/1% AA. The sample was centrifuged at 1000 rpm for 5 min (C). After centrifugation, supernatant was removed and 1 ml of DMEM/2% FBS/1% AA was added. Then, 10 ml of RBC lysis buffer was added and gently pipetted several times. After pipetting, the sample was incubated at 4°C for 12 min (D). After incubation, a sample was gently pipetted several times and centifuged at 1000 rpm for 5 min again. Supernatant was removed, washed with 5 ml of PBS and centrifuged at 1000 rpm for 5 min. This process was done several times to remove RBC until the supernatant looked transparent (E).

**Supplemental Figure 2. Proliferative capacity of primary and freeze-thawed MenSCs from infertile patients**

(A) Phase-contrast photomicrographs of primary and freeze-thawed MenSCs from infertile patients at different time points. (B) MenSCs (1.0 × 10^5^) at passage 4-7 were plated into 6-well plates with DMEM/10% FBS (n=3 in each group). Cells were counted at day 1, 3, 5 and 7. Proliferative capacity was not significantly decreased until day 7 even after the freeze-thawed procedure. Statistical significance is shown as *P <0.05 and **P <0.01. ‘ns’ means ‘not significant’. Black bars are 500 μm.

**Supplemental Figure 3. Proteomics analysis of CM from volunteer- and infertile patient-derived MenSCs**

(A) Quantitative scatterplot indicated that proteins contained in CM from volunteer- and infertile patient-derived MenSCs were similar to each other. (B) Number of ‘unique proteins’ was similar in the volunteer-derived and infertile patient-derived CM in the categories of ‘biological process’, ‘cellular component’ and ‘molecular function’. (C) Venn diagram showing that total 51 proteins were common between volunteer- and infertile patient-derived CM. (D) The 51 proteins were primarily categorized into subgroups of ‘biological process’ such as ‘cellular process’, ‘biological regulation’, ‘developmental process’, ‘multicellular organismal process’, ‘response to stimulus’, ‘localization’, and ‘biological adhesion’.

**Supplemental Figure 4. Systemic biodistribution of intrauterine transplanted MenSCs in major organs**.

Histopathological findings of major organs at day 1, 7 and 30 after intrauterine transplantation of MenSCs (n=3 in each group). There were no human vimentin-positive cells and tumor formation in brain, heart, lungs, liver, spleen, pancreas, kidneys, adrenal glands and ovaries. Black bars are 500 μm.

## Supplemental Tables

**Supplemental Table 1**. List of antibodies for immunohistochemistry and flow-cytometry

**Supplemental Table 2**. List of primer sequences for quantitative reverse transcription polymerase chain reaction

**Supplemental Table 3**. Patient characteristics of samples with unsuccessful primary culture of menstrual-blood derived cells

**Supplemental Table 4**. Flow cytometric analysis for mesenchymal stem cell markers

**Supplemental Table 5**. List of common proteins in the CM from volunteer and infertile patient menstrual blood-derived cells

**Supplemental Table 6**. Body and organ weights of the mice in each group Supplemental Table 7. Blood sample test of the mice in each group

## Supplemental File

**Supplemental File 1**. Movie of intrauterine curettage

