## Supplementary figures and images for "Novel therapeutic strategies for injured endometrium: Autologous intrauterine transplantation of menstrual blood-derived cells from infertile patients"

### Supplemental Figure 1

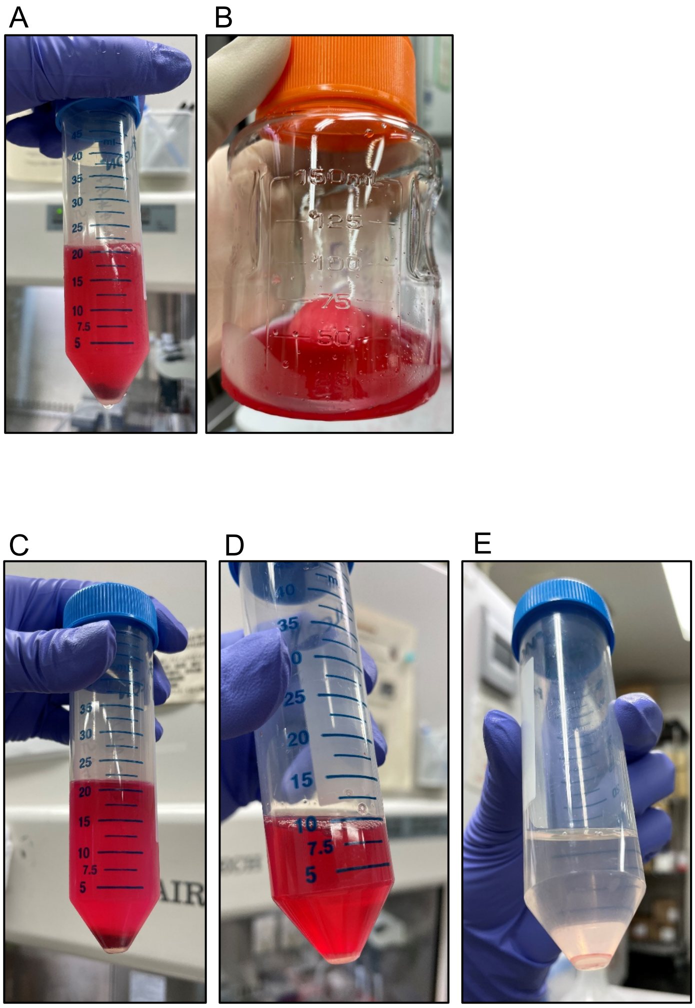


Supplemental figure 1.

### Supplemental Figure 2

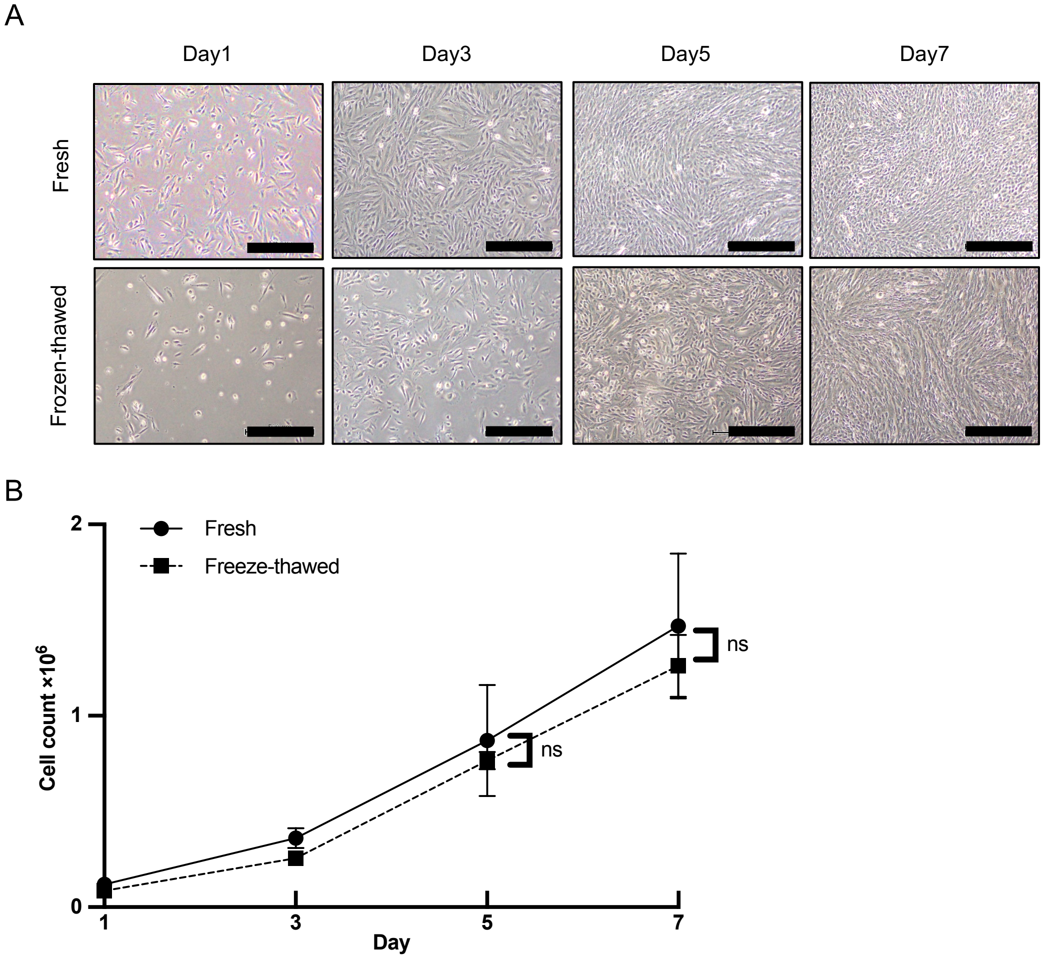


Supplemental figure 2.

### Supplemental Figure 3

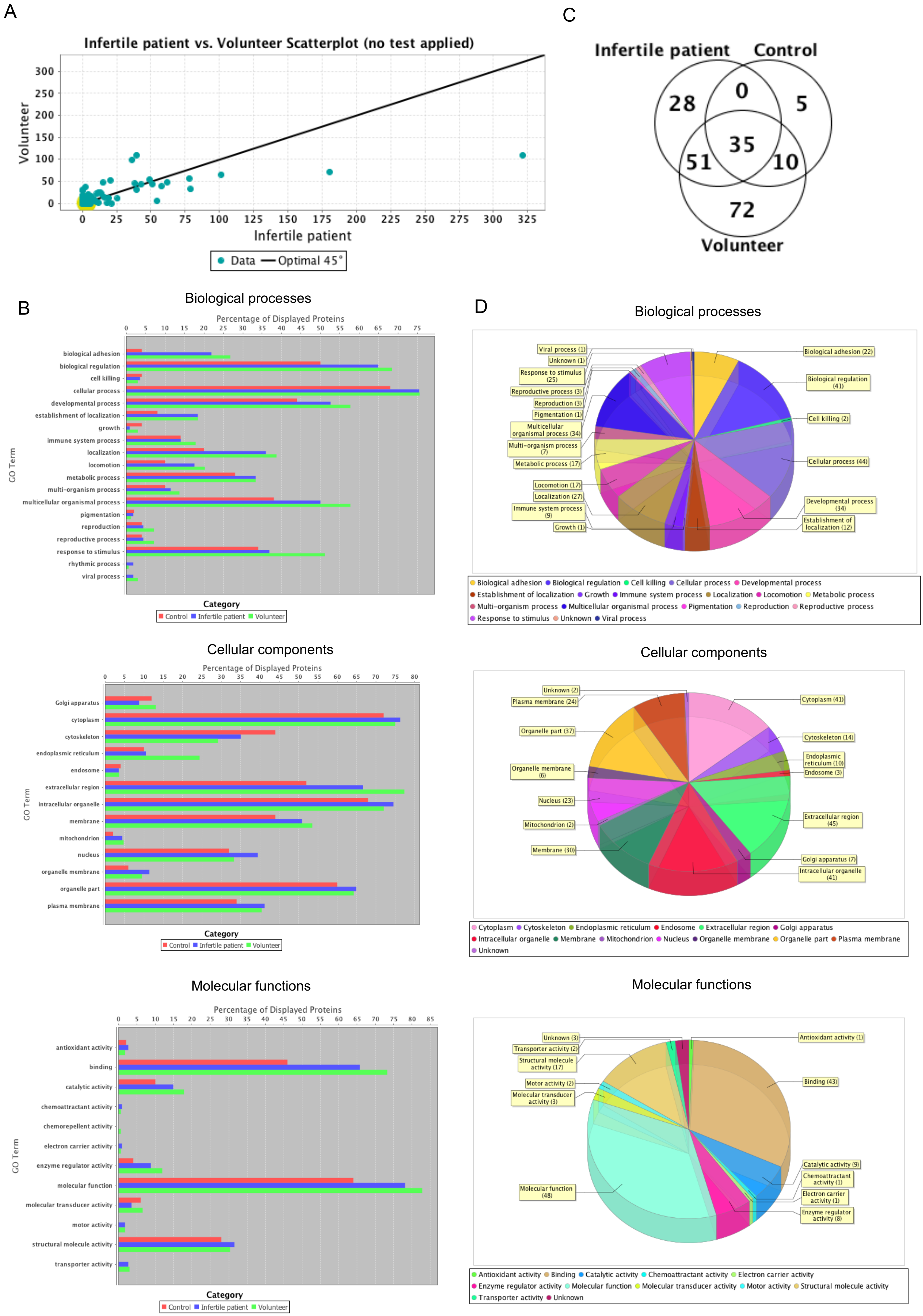


Supplemental Figure 3.

### Supplemental Figure 4

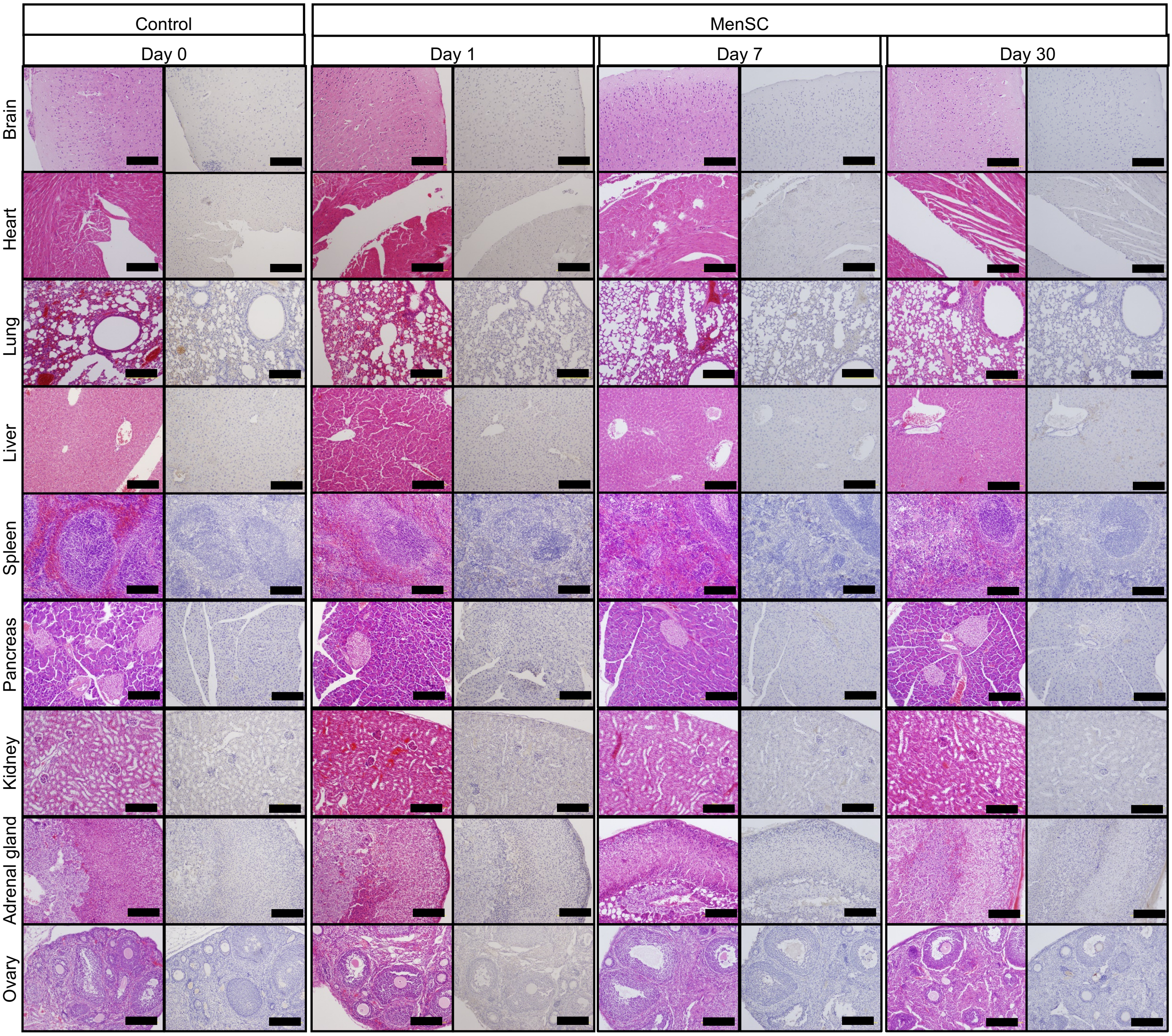


Supplemental figure 4.
