## Supplemental Table 1 for "Novel therapeutic strategies for injured endometrium: Autologous intrauterine transplantation of menstrual blood-derived cells from infertile patients"

**Supplemental Table 1.** List of antibodies for immunohistochemistry and flow-cytometry

| Name | Clone | Species reactivity | Host | Company | Dilution |
|---|---|---|---|---|---|
| Immunohistochemistry | | | | | |
| Primary antibodies | | | | | |
| Anti-CD34 antibody | Ab81289 | Mouse, rat, human | Rabbit, Monoclonal IgG | Abcam | 1/2500 |
| Anti-Ki-67 antibody | Ab15581 | Mouse, human | Rabbit, Polyclonal IgG | Abcam | 1/300 |
| Anti-human Vimentin antibody | M7020 | human | Mouse, Monoclonal IgG | Dako | 1/100 |
| Secondary antibodies | | | | | |
| -Cellstain®- DAPI solution | D523 | None | | DOJINDO | 1/1000 |
| Rabbit anti-goat IgG Secondary antibody, Alexa Fluor 488 | A11078 | None | | Invitrogen | 1/500 |
| Goat anti-mouse IgG1 Secondary antibody, Alexa Fluor 546 | A21123 | None | | Invitrogen | 1/500 |
| Flow-cytometry | | | | | |
| Anti-CD73 Antibody, anti-human, REAfinity™ | AD2 | human | human cell line, monoclonal IgG1 | Miltenyi Biotec Inc | 1/50 |
| Anti-CD90 Antibody, anti-human, REAfinity™ | DG3 | human | human cell line, monoclonal IgG1 | Miltenyi Biotec Inc | 1/50 |
| Anti-CD105 Antibody, anti-human, REAfinity™ | 43A4E1 | human | human cell line, monoclonal IgG1 | Miltenyi Biotec Inc | 1/50 |
| Anti-CD14 Antibody, anti-human, REAfinity™ | REA599 | human | human cell line, monoclonal IgG1 | Miltenyi Biotec Inc | 1/50 |
| Anti-CD19 Antibody, anti-human, REAfinity™ | REA675 | human | human cell line, monoclonal IgG1 | Miltenyi Biotec Inc | 1/50 |
| Anti-CD34 Antibody, anti-human, REAfinity™ | REA1164 | human | human cell line, monoclonal IgG1 | Miltenyi Biotec Inc | 1/50 |
| Anti-CD45 Antibody, anti-human, REAfinity™ | REA747 | human | human cell line, monoclonal IgG1 | Miltenyi Biotec Inc | 1/50 |
| Anti-HLA-DR Antibody, anti-human, REAfinity™ | REA805 | human | human cell line, monoclonal IgG1 | Miltenyi Biotec Inc | 1/50 |
| Isotype Control Antibody mouse IgG1, PE | IS5-21F5 | human | mouse, monoclonal IgG1 | Miltenyi Biotec Inc | 1/50 |
| REA Control Antibody(S), human IgG1, APC, REAfinity | REA293 | human | human cell line, monoclonal IgG1 | Miltenyi Biotec Inc | 1/50 |

DAPI, 4',6-Diamidino-2-Phenylindole, dihydrochloride.
