## Supplemental Table 2 for "Novel therapeutic strategies for injured endometrium: Autologous intrauterine transplantation of menstrual blood-derived cells from infertile patients"

**Supplemental Table 2.** List of primer sequences for quantitative reverse transcription polymerase chain reaction

| Gene | Primer sequence | 5' to 3' |
|---|---|---|
| *VEGFA* | Forward | 5' TCAGTTCGAGGAAAGGGAAA 3' |
| | Reverse | 5' GCGAGTCTGTGTTTTTGCAG 3' |
| *FGF-1* | Forward | 5' GGACACCGAAGGGCTTTTAT 3' |
| | Reverse | 5' ACAGCTCCCGTTCTTCTTGA 3' |
| *FGF-2* | Forward | 5' CCTTGCTATGAAGGAAGATGG 3' |
| | Reverse | 5' CCGTTTTGGATCCGAGTTTA 3' |
| *EGF* | Forward | 5' CGTGTGCATGCATATTGAATC 3' |
| | Reverse | 5' TTCCCACCATCGTAGGTCTC 3' |
| *GAPDH* | Forward | 5' TGTTGCCATCAATGACCCCTT 3' |
| | Reverse | 5' CTCCACGACGTACTCAGCG 3' |

VEGFA, vascular endothelial growth factor A; FGF, fibroblast growth factor; EGF, epidermal growth factor; GAPDH, glyceraldehyde 3-phosphate dehydrogenase.
