## Supplemental Table 3 for "Novel therapeutic strategies for injured endometrium: Autologous intrauterine transplantation of menstrual blood-derived cells from infertile patients"

**Supplemental Table 3.** Patient characteristics of samples with unsuccessful primary culture of menstrual blood-derived cells

| Sample # | Age, y | Gravida | Parity | BMI, $kg/m^2$ | Cause for infertility | Duration of infertility, y | Treatment method | Menstrual day | Collection method |
|---|---|---|---|---|---|---|---|---|---|
| 5 | 48 | 1 | 1 | 23.3 | Ovarian insufficiency | 0.5 | TI | D3 | Cotton |
| 7 | 37 | 2 | 1 | 21.1 | PCOS | 1.1 | TI | D2 | Cotton |
| 11 | 43 | 4 | 1 | 19.6 | Endometriosis | 0.8 | ART | D5 | Cotton |
| 12 | 35 | 0 | 0 | 19.7 | Unexplained | 0.4 | TI | D3 | Cotton |
| 15 | 30 | 0 | 0 | 20.1 | PCOS | 0.6 | IUI | D4 | Syringe |
| 16 | 37 | 0 | 0 | 18.1 | PCOS | 0.7 | TI | D3 | Cotton |

y, year; BMI, body mass index; PCOS, polycystic ovary syndrome; TI, timed intercourse; ART, assisted reproductive technology; IUI, intrauterine insemination; D, day.
