## Supplemental Table 4 for "Novel therapeutic strategies for injured endometrium: Autologous intrauterine transplantation of menstrual blood-derived cells from infertile patients"

**Supplemental Table 4.** Flow cytometric analysis for mesenchymal stem cell markers

| Antigen | Volunteers (%, mean ± SD) | Infertile patients (%, mean ± SD) | *P*-value |
|---|---|---|---|
| CD73 | 96.07 ± 4.31 | 98.80 ± 0.36 | 0.33 |
| CD90 | 85.57 ± 6.93 | 75.83 ± 11.93 | 0.48 |
| CD105 | 94.63 ± 4.14 | 96.63 ± 3.50 | 0.66 |
| CD14 | 0.70 ± 0.63 | 1.03 ± 0.25 | 0.49 |
| CD19 | 0.73 ± 0.61 | 0.98 ± 0.21 | 0.57 |
| CD34 | 0.70 ± 0.64 | 1.26 ± 0.54 | 0.44 |
| CD45 | 0.83 ± 0.42 | 1.45 ± 0.28 | 0.16 |
| HLA-DR | 0.65 ± 0.54 | 0.89 ± 0.04 | 0.48 |

These measurements were conducted entirely independent of all other variables (n=3).
