## Supplemental Table 5 for "Novel therapeutic strategies for injured endometrium: Autologous intrauterine transplantation of menstrual blood-derived cells from infertile patients"

**Supplemental Table 5.** List of proteins common to CM from volunteer-derived and infertile patient-derived menstrual blood derived cells

| # | Proteins | Accession Number | biological adhesion | biological regulation |
|---|---|---|---|---|
| 1 | Thrombospondin-1 OS=Homo sapiens OX=9606 GN=THBS1 PE=1 SV=2 | P07996 | cell adhesion | negative regulation of angiogenesis, negative regulation of antigen processing and presentation of peptide or polysaccharide antigen via MHC class II, negative regulation of apoptotic process, negative regulation of blood vessel endothelial cell proliferation involved in sprouting angiogenesis, negative regulation of cGMP-mediated signaling, negative regulation of cell migration involved in sprouting angiogenesis, negative regulation of cell proliferation, negative regulation of cell-matrix adhesion, negative regulation of cysteine-type endopeptidase activity involved in apoptotic process, negative regulation of dendritic cell antigen processing and presentation, negative regulation of endothelial cell chemotaxis, negative regulation of endothelial cell proliferation, negative regulation of extrinsic apoptotic signaling pathway, negative regulation of fibrinolysis, negative regulation of fibroblast growth factor receptor signaling pathway, negative regulation of focal adhesion assembly, negative regulation of interleukin-12 production, negative regulation of nitric oxide mediated signal transduction, negative regulation of plasma membrane long-chain fatty acid transport, negative regulation of plasminogen activation, negative regulation of sprouting angiogenesis, positive regulation of MAP kinase activity, positive regulation of angiogenesis, positive regulation of blood vessel endothelial cell migration, positive regulation of cell migration, positive regulation of chemotaxis, positive regulation of endothelial cell apoptotic process, positive regulation of extrinsic apoptotic signaling pathway via death domain receptors, positive regulation of fibroblast migration, positive regulation of macrophage activation, positive regulation of macrophage chemotaxis, positive regulation of phosphorylation, positive regulation of protein kinase B signaling, positive regulation of reactive oxygen species metabolic process, positive regulation of smooth muscle cell proliferation, positive regulation of transforming growth factor beta receptor signaling pathway, positive regulation of transforming growth factor beta1 production, positive regulation of translation, positive regulation of tumor necrosis factor production |
| 2 | Fibronectin OS=Homo sapiens OX=9606 GN=FN1 PE=1 SV=5 | P02751 | calcium-independent cell-matrix adhesion, cell-matrix adhesion, platelet aggregation, substrate adhesion-dependent cell spreading | blood coagulation, fibrin clot formation, integrin-mediated signaling pathway, negative regulation of transforming growth factor beta production, platelet aggregation, positive regulation of axon extension, positive regulation of cell proliferation, positive regulation of fibroblast proliferation, positive regulation of gene expression, positive regulation of phosphatidylinositol 3-kinase signaling, positive regulation of substrate-dependent cell migration, cell attachment to substrate, regulation of ERK1 and ERK2 cascade, regulation of cell shape, regulation of protein phosphorylation |
| 3 | Collagen alpha-1(XII) chain OS=Homo sapiens OX=9606 GN=COL12A1 PE=1 SV=2 | Q99715 | cell adhesion | |
| 4 | Latent-transforming growth factor beta-binding protein 2 OS=Homo sapiens OX=9606 GN=LTBP2 PE=1 SV=3 | Q14767 | | transforming growth factor beta receptor signaling pathway |
| 5 | Thrombospondin-2 OS=Homo sapiens OX=9606 GN=THBS2 PE=1 SV=2 | P35442 | cell adhesion | negative regulation of angiogenesis, positive regulation of synapse assembly |
| 6 | Agrin OS=Homo sapiens OX=9606 GN=AGRN PE=1 SV=6 | O00468 | | G-protein coupled acetylcholine receptor signaling pathway, positive regulation of GTPase activity, positive regulation of filopodium assembly, positive regulation of synaptic growth at neuromuscular junction, positive regulation of transcription from RNA polymerase II promoter, signal transduction |
| 7 | Cluster of Actin, cytoplasmic 1 OS=Homo sapiens OX=9606 GN=ACTB PE=1 SV=1 (P60709) | P60709 [5] | platelet aggregation | maintenance of permeability of blood-brain barrier, negative regulation of apoptotic process, negative regulation of cell differentiation, negative regulation of protein binding, platelet aggregation, positive regulation of T cell differentiation, positive regulation of cell differentiation, positive regulation of cell proliferation, positive regulation of double-strand break repair, positive regulation of double-strand break repair via homologous recombination, positive regulation of gene expression, positive regulation of myoblast differentiation, positive regulation of norepinephrine uptake, positive regulation of stem cell population maintenance, positive regulation of transcription, DNA-templated, regulation of G0 to G1 transition, regulation of G1/S transition of mitotic cell cycle, regulation of cyclin-dependent protein serine/threonine kinase activity, regulation of double-strand break repair, regulation of mitotic metaphase/anaphase transition, regulation of norepinephrine uptake, regulation of nucleotide-excision repair, regulation of protein localization to plasma membrane, regulation of transcription from RNA polymerase II promoter, regulation of transmembrane transporter activity, retina homeostasis |
| 8 | Nidogen-1 OS=Homo sapiens OX=9606 GN=NID1 PE=1 SV=3 | P14543 | cell-matrix adhesion | positive regulation of cell adhesion, positive regulation of cell-substrate adhesion, positive regulation of integrin-mediated signaling pathway, positive regulation of muscle cell differentiation |
| 9 | Cluster of Alpha-actinin-4 OS=Homo sapiens OX=9606 GN=ACTN4 PE=1 SV=2 (O43707) | O43707 [2] | focal adhesion assembly | negative regulation of cellular component movement, negative regulation of substrate adhesion-dependent cell spreading, peroxisome proliferator activated receptor signaling pathway, positive regulation of NIK/NF-kappaB signaling, positive regulation of cell migration, positive regulation of cellular component movement, positive regulation of sodium:proton antiporter activity, regulation of apoptotic process, regulation of nucleic acid-templated transcription, retinoic acid receptor signaling pathway, tumor necrosis factor-mediated signaling pathway |
| 10 | Lysyl oxidase homolog 2 OS=Homo sapiens OX=9606 GN=LOXL2 PE=1 SV=1 | Q9Y4K0 | cell adhesion | negative regulation of stem cell population maintenance, negative regulation of transcription from RNA polymerase II promoter, negative regulation of transcription, DNA-templated, positive regulation of chondrocyte differentiation, positive regulation of epithelial to mesenchymal transition |
| 11 | Glypican-1 OS=Homo sapiens OX=9606 GN=GPC1 PE=1 SV=2 | P35052 | | negative regulation of fibroblast growth factor receptor signaling pathway, positive regulation of skeletal muscle cell differentiation, regulation of protein localization to membrane |
| 12 | Nidogen-2 OS=Homo sapiens OX=9606 GN=NID2 PE=1 SV=3 | Q14112 | cell-matrix adhesion | |
| 13 | Junction plakoglobin OS=Homo sapiens OX=9606 GN=JUP PE=1 SV=3 | P14923 | bundle of His cell-Purkinje myocyte adhesion involved in cell communication, cell-cell adhesion, endothelial cell-cell adhesion | bundle of His cell-Purkinje myocyte adhesion involved in cell communication, negative regulation of blood vessel endothelial cell migration, positive regulation of angiogenesis, positive regulation of canonical Wnt signaling pathway, positive regulation of cell-matrix adhesion, positive regulation of protein import into nucleus, positive regulation of sequence-specific DNA binding transcription factor activity, positive regulation of transcription from RNA polymerase II promoter, regulation of cell proliferation, regulation of heart rate by cardiac conduction, regulation of ventricular cardiac muscle cell action potential |
| 14 | Amyloid-beta precursor protein OS=Homo sapiens OX=9606 GN=APP PE=1 SV=3 | P05067 | cell adhesion | Notch signaling pathway, adenylate cyclase-activating G-protein coupled receptor signaling pathway, adenylate cyclase-inhibiting G-protein coupled receptor signaling pathway, calcium-mediated signaling, cellular copper ion homeostasis, ionotropic glutamate receptor signaling pathway, modulation of age-related behavioral decline, negative regulation of blood circulation, negative regulation of canonical Wnt signaling pathway, negative regulation of cell proliferation, negative regulation of dendritic spine maintenance, negative regulation of gene expression, negative regulation of long-term synaptic potentiation, negative regulation of mitochondrion organization, negative regulation of neuron death, negative regulation of neuron differentiation, negative regulation of pri-miRNA transcription from RNA polymerase II promoter, negative regulation of protein localization to nucleus, negative regulation of transcription from RNA polymerase II promoter, positive regulation of ERK1 and ERK2 cascade, positive regulation of G-protein coupled receptor internalization, positive regulation of G-protein coupled receptor protein signaling pathway, positive regulation of G2/M transition of mitotic cell cycle, positive regulation of JNK cascade, positive regulation of MAP kinase activity, positive regulation of MAPK cascade, positive regulation of NF-kappaB transcription factor activity, positive regulation of NIK/NF-kappaB signaling, positive regulation of T cell migration, positive regulation of amyloid fibril formation, positive regulation of apoptotic process, positive regulation of aspartic-type endopeptidase activity involved in amyloid precursor protein catabolic process, positive regulation of cell activation, positive regulation of cellular response to thapsigargin, positive regulation of cellular response to tunicamycin, positive regulation of chemokine production, positive regulation of cysteine-type endopeptidase activity involved in apoptotic process, positive regulation of cytosolic calcium ion concentration, positive regulation of endothelin secretion, positive regulation of excitatory postsynaptic potential, positive regulation of gene expression, positive regulation of glycolytic process, positive regulation of histone acetylation, positive regulation of inflammatory response, positive regulation of interferon-gamma production, positive regulation of interleukin-1 beta production, positive regulation of interleukin-6 production, positive regulation of long term synaptic depression, positive regulation of long-term synaptic potentiation, positive regulation of membrane protein ectodomain proteolysis, positive regulation of mitotic cell cycle, positive regulation of monocyte chemotaxis, positive regulation of neuron apoptotic process, positive regulation of neuron death, positive regulation of neuron differentiation, positive regulation of nitric oxide biosynthetic process, positive regulation of oxidative stress-induced neuron death, positive regulation of peptidyl-serine phosphorylation, positive regulation of peptidyl-threonine phosphorylation, positive regulation of phosphorylation, positive regulation of protein binding, positive regulation of protein import, positive regulation of protein kinase A signaling, positive regulation of protein kinase B signaling, positive regulation of protein tyrosine kinase activity, positive regulation of receptor binding, positive regulation of response to endoplasmic reticulum stress, positive regulation of superoxide anion generation, positive regulation of tau-protein kinase activity, positive regulation of transcription from RNA polymerase II promoter, positive regulation of tumor necrosis factor production, regulation of MAPK cascade, regulation of NMDA receptor activity, regulation of Wnt signaling pathway, regulation of acetylcholine-gated cation channel activity, regulation of amyloid fibril formation, regulation of beta-amyloid clearance, regulation of dendritic spine maintenance, regulation of epidermal growth factor-activated receptor activity, regulation of long-term neuronal synaptic plasticity, regulation of multicellular organism growth, regulation of peptidyl-tyrosine phosphorylation, regulation of presynapse assembly, regulation of protein tyrosine kinase activity, regulation of response to calcium ion, regulation of synapse structure or activity, regulation of toll-like receptor signaling pathway, regulation of transcription from RNA polymerase II promoter, regulation of translation, smooth endoplasmic reticulum calcium ion homeostasis |
| 15 | Laminin subunit gamma-1 OS=Homo sapiens OX=9606 GN=LAMC1 PE=1 SV=3 | P11047 | substrate adhesion-dependent cell spreading | maintenance of permeability of blood-brain barrier, positive regulation of cell adhesion, positive regulation of epithelial cell proliferation, positive regulation of integrin-mediated signaling pathway, positive regulation of muscle cell differentiation |
| 16 | Galectin-3-binding protein OS=Homo sapiens OX=9606 GN=LGALS3BP PE=1 SV=1 | Q08380 | cell adhesion | signal transduction |
| 17 | Matrix-remodeling-associated protein 5 OS=Homo sapiens OX=9606 GN=MXRA5 PE=1 SV=3 | Q9NR99 | | |
| 18 | Testican-1 OS=Homo sapiens OX=9606 GN=SPOCK1 PE=1 SV=1 | Q08629 | cell adhesion | negative regulation of cell-substrate adhesion, negative regulation of endopeptidase activity, negative regulation of neuron projection development, regulation of cell growth |
| 19 | Laminin subunit beta-1 OS=Homo sapiens OX=9606 GN=LAMB1 PE=1 SV=2 | P07942 | neuronal-glial interaction involved in cerebral cortex radial glia guided migration, substrate adhesion-dependent cell spreading | positive regulation of cell adhesion, positive regulation of cell migration, positive regulation of epithelial cell proliferation, positive regulation of integrin-mediated signaling pathway, positive regulation of muscle cell differentiation |
| 20 | Adipocyte enhancer-binding protein 1 OS=Homo sapiens OX=9606 GN=AEBP1 PE=1 SV=1 | Q8IUX7 | | negative regulation of transcription from RNA polymerase II promoter, regulation of collagen fibril organization |
| 21 | Decorin OS=Homo sapiens OX=9606 GN=DCN PE=1 SV=1 | P07585 | | negative regulation of angiogenesis, negative regulation of endothelial cell migration, negative regulation of vascular endothelial growth factor signaling pathway, positive regulation of autophagy, positive regulation of macroautophagy, positive regulation of mitochondrial depolarization, positive regulation of mitochondrial fission, positive regulation of phosphatidylinositol 3-kinase signaling, positive regulation of transcription from RNA polymerase II promoter |
| 22 | Desmoglein-1 OS=Homo sapiens OX=9606 GN=DSG1 PE=1 SV=2 | Q02413 | calcium-dependent cell-cell adhesion via plasma membrane cell adhesion molecules, homophilic cell adhesion via plasma membrane adhesion molecules | protein stabilization |
| 23 | Endosialin OS=Homo sapiens OX=9606 GN=CD248 PE=1 SV=1 | Q9HCU0 | | positive regulation of cell proliferation, positive regulation of endothelial cell apoptotic process |
| 24 | Versican core protein OS=Homo sapiens OX=9606 GN=VCAN PE=1 SV=3 | P13611 | cell adhesion | |
| 25 | Myosin-9 OS=Homo sapiens OX=9606 GN=MYH9 PE=1 SV=4 | P35579 | platelet aggregation | cortical granule exocytosis, integrin-mediated signaling pathway, negative regulation of actin filament severing, platelet aggregation, positive regulation of protein processing in phagocytic vesicle, regulation of cell shape, regulation of plasma membrane repair |
| 26 | C-type lectin domain family 11 member A OS=Homo sapiens OX=9606 GN=CLEC11A PE=1 SV=1 | Q9Y240 | | positive regulation of cell proliferation |
| 27 | Elongation factor 1-alpha 1 OS=Homo sapiens OX=9606 GN=EEF1A1 PE=1 SV=1 | P68104 (+1) | | regulation of D-erythro-sphingosine kinase activity, regulation of chaperone-mediated autophagy |
| 28 | Glyceraldehyde-3-phosphate dehydrogenase OS=Homo sapiens OX=9606 GN=GAPDH PE=1 SV=3 | P04406 | | killing by host of symbiont cells, killing of cells of other organism, negative regulation of endopeptidase activity, negative regulation of translation, positive regulation of I-kappaB kinase/NF-kappaB signaling, positive regulation of type I interferon production, protein stabilization, regulation of macroautophagy |
| 29 | Pyruvate kinase PKM OS=Homo sapiens OX=9606 GN=PKM PE=1 SV=4 | P14618 | | positive regulation of cytoplasmic translation, positive regulation of sprouting angiogenesis |
| 30 | Biglycan OS=Homo sapiens OX=9606 GN=BGN PE=1 SV=2 | P21810 | | |
| 31 | Tubulin alpha-1B chain OS=Homo sapiens OX=9606 GN=TUBA1B PE=1 SV=1 | P68363 (+2) | | |
| 32 | Sushi repeat-containing protein SRPX2 OS=Homo sapiens OX=9606 GN=SRPX2 PE=1 SV=1 | O60687 | cell-cell adhesion | positive regulation of cell migration involved in sprouting angiogenesis, positive regulation of synapse assembly, regulation of phosphorylation |
| 33 | Peroxiredoxin-1 OS=Homo sapiens OX=9606 GN=PRDX1 PE=1 SV=1 | Q06830 | | cell redox homeostasis, erythrocyte homeostasis, regulation of NIK/NF-kappaB signaling, regulation of stress-activated MAPK cascade, retina homeostasis |
| 34 | Annexin A2 OS=Homo sapiens OX=9606 GN=ANXA2 PE=1 SV=2 | P07355 | | negative regulation of low-density lipoprotein particle receptor catabolic process, negative regulation of receptor internalization, positive regulation of exocytosis, positive regulation of low-density lipoprotein particle clearance, positive regulation of low-density lipoprotein particle receptor binding, positive regulation of low-density lipoprotein receptor activity, positive regulation of plasma membrane repair, positive regulation of plasminogen activation, positive regulation of receptor recycling, positive regulation of receptor-mediated endocytosis involved in cholesterol transport, positive regulation of vacuole organization, positive regulation of vesicle fusion |
| 35 | Protein S100-A8 OS=Homo sapiens OX=9606 GN=S100A8 PE=1 SV=1 | P05109 | neutrophil aggregation | activation of cysteine-type endopeptidase activity involved in apoptotic process, positive regulation of NF-kappaB transcription factor activity, positive regulation of cell growth, positive regulation of inflammatory response, positive regulation of intrinsic apoptotic signaling pathway, positive regulation of peptide secretion, regulation of cytoskeleton organization, sequestering of zinc ion |
| 36 | 14-3-3 protein zeta/delta OS=Homo sapiens OX=9606 GN=YWHAZ PE=1 SV=1 | P63104 | | ERK1 and ERK2 cascade, negative regulation of apoptotic process, negative regulation of transcription from RNA polymerase II promoter, regulation of ERK1 and ERK2 cascade, regulation of synapse maturation, signal transduction |
| 37 | Histone H3.1 OS=Homo sapiens OX=9606 GN=H3C1 PE=1 SV=2 | P68431 (+3) | | regulation of gene expression, epigenetic |
| 38 | Histone H2A type 1-B/E OS=Homo sapiens OX=9606 GN=H2AC4 PE=1 SV=2 | P04908 (+13) | | negative regulation of cell proliferation |
| 39 | Histone H2B type 2-K1 OS=Homo sapiens OX=9606 GN=H2BK1 PE=3 SV=1 | A0A2R8Y619 (+15) | | |
| 40 | Biogenesis of lysosome-related organelles complex 1 subunit 3 OS=Homo sapiens OX=9606 GN=BLOC1S3 PE=1 SV=1 | Q6QNY0 | | platelet activation, positive regulation of natural killer cell activation |
| 41 | Filaggrin-2 OS=Homo sapiens OX=9606 GN=FLG2 PE=1 SV=1 | Q5D862 | cell adhesion | establishment of skin barrier |
| 42 | Skin-specific protein 32 OS=Homo sapiens OX=9606 GN=XP32 PE=1 SV=1 | Q5T750 | | |
| 43 | Serglycin OS=Homo sapiens OX=9606 GN=SRGN PE=1 SV=3 | P10124 | | maintenance of granzyme B location in T cell secretory granule, maintenance of protease location in mast cell secretory granule, modulation of synaptic transmission, negative regulation of bone mineralization, negative regulation of cytokine production |
| 44 | Amyloid beta precursor like protein 2 OS=Homo sapiens OX=9606 GN=APLP2 PE=1 SV=2 | Q06481 | | G-protein coupled receptor signaling pathway |
| 45 | Calmodulin-like protein 5 OS=Homo sapiens OX=9606 GN=CALML5 PE=1 SV=2 | Q9NZT1 | | signal transduction |
| 46 | Low-density lipoprotein receptor-related protein 8 OS=Homo sapiens OX=9606 GN=LRP8 PE=1 SV=4 | Q14114 | | cytokine-mediated signaling pathway, modulation of synaptic transmission, positive regulation of CREB transcription factor activity, positive regulation of dendrite development, positive regulation of dendritic spine morphogenesis, positive regulation of peptidyl-tyrosine phosphorylation, positive regulation of protein tyrosine kinase activity, reelin-mediated signaling pathway, regulation of apoptotic process, regulation of innate immune response, signal transduction |
| 47 | Peroxisomal multifunctional enzyme type 2 OS=Homo sapiens OX=9606 GN=HSD17B4 PE=1 SV=3 | P51659 | | |
| 48 | Heat shock protein beta-1 OS=Homo sapiens OX=9606 GN=HSPB1 PE=1 SV=2 | P04792 | platelet aggregation | intracellular signal transduction, negative regulation of apoptotic process, negative regulation of oxidative stress-induced intrinsic apoptotic signaling pathway, negative regulation of protein kinase activity, platelet aggregation, positive regulation of angiogenesis, positive regulation of blood vessel endothelial cell migration, positive regulation of endothelial cell chemotaxis, positive regulation of endothelial cell chemotaxis by VEGF-activated vascular endothelial growth factor receptor signaling pathway, positive regulation of interleukin-1 beta production, positive regulation of tumor necrosis factor production, regulation of I-kappaB kinase/NF-kappaB signaling, regulation of autophagy, regulation of translational initiation, retina homeostasis |
