## Supplemental Table 6 for "Novel therapeutic strategies for injured endometrium: Autologous intrauterine transplantation of menstrual blood-derived cells from infertile patients"

**Supplemental Table 6.** Relative body and organ weights of the mice in each group

| Variables, g, n=4 | Sham | Injured | MenSCs | P value |
|---|---|---|---|---|
| Body weight | 27.70 ± 2.37 | 28.11 ± 2.03 | 27.29 ± 0.93 | 0.82 |
| Body weight change | -0.18 ± 0.88 | 0.23 ± 0.92 | 0.52 ± 0.52 | 0.48 |
| Brain | 0.489 ± 0.045 | 0.523 ± 0.029 | 0.479 ± 0.039 | 0.28 |
| Heart | 0.149 ± 0.016 | 0.154 ± 0.004 | 0.144 ± 0.014 | 0.51 |
| Lung | 0.204 ± 0.044 | 0.205 ± 0.038 | 0.194 ± 0.003 | 0.99 |
| Liver | 1.89 ± 0.11 | 1.92 ± 0.19 | 1.98 ± 0.17 | 0.74 |
| Spleen | 0.120 ± 0.024 | 0.122 ± 0.023 | 0.116 ± 0.028 | 0.95 |
| Pancreas | 0.242 ± 0.061 | 0.221 ± 0.042 | 0.235 ± 0.064 | 0.86 |
| Kidney | 0.474 ± 0.041 | 0.488 ± 0.039 | 0.486 ± 0.040 | 0.86 |
| Adrenal gland | 0.014 ± 0.003 | 0.009 ± 0.002 | 0.010 ± 0.003 | 0.09 |
| Ovary | 0.019 ± 0.005 | 0.028 ± 0.004 | 0.023 ± 0.007 | 0.13 |

Data are presented as mean ± SD. One-way ANOVA test was conducted for calculating statistical difference.
