## Supplemental Table 7 for "Novel therapeutic strategies for injured endometrium: Autologous intrauterine transplantation of menstrual blood-derived cells from infertile patients"

**Supplemental Table 7.** Blood sample test of the mice in each group

| Variables, n=4 | Sham | Injured | MenSCs | P value |
|---|---|---|---|---|
| WBC (×10^2/μL) | 47.50 ± 14.53 | 45.75 ± 11.44 | 27.75 ± 9.39 | 0.07 |
| RBC (×10^4/μL) | 825.25 ± 66.83 | 801.75 ± 82.27 | 767.50 ± 14.98 | 0.44 |
| Hb (g/dL) | 13.32 ± 1.07 | 13.00 ± 1.43 | 12.43 ± 0.40 | 0.50 |
| HCT (%) | 40.45 ± 3.78 | 39.18 ± 3.58 | 37.13 ± 1.41 | 0.35 |
| MCV (fL) | 48.9 ± 0.96 | 48.93 ± 1.74 | 48.38 ± 1.06 | 0.80 |
| MCH (pg) | 16.15 ± 0.12 | 16.20 ± 0.30 | 16.18 ± 0.16 | 0.98 |
| MCHC (g/dL) | 33.00 ± 0.65 | 33.15 ± 0.57 | 33.45 ± 0.24 | 0.48 |
| PLT (×10^4/μL) | 111.85 ± 20.81 | 108.75 ± 6.40 | 121.25 ± 4.03 | 0.92 |
| TP (g/dL) | 4.98 ± 0.33 | 4.98 ± 0.26 | 4.73 ± 0.30 | 0.42 |
| A/G ratio | 1.68 ± 0.21 | 1.60 ± 0.22 | 1.83 ± 0.05 | 0.23 |
| Alb (g/dL) | 3.10 ± 0.14 | 3.03 ± 0.10 | 3.05 ± 0.17 | 0.75 |
| BUN (mg/dL) | 26.53 ± 4.23 | 22.80 ± 1.24 | 23.48 ± 2.09 | 0.19 |
| Cre (mg/dL) | 0.11 ± 0.02 | 0.09 ± 0.01 | 0.10 ± 0.01 | 0.45 |
| Na (mEq/L) | 148.00 ± 4.08 | 149.75 ± 2.36 | 148.25 ± 4.35 | 0.77 |
| Cl (mEq/L) | 108.75 ± 2.63 | 109.75 ± 1.89 | 108.00 ± 1.83 | 0.53 |
| ALT (IU/L) | 21.00 ± 4.97 | 21.00 ± 5.03 | 27.75 ± 5.38 | 0.19 |
| ALP (IU/L) | 248.25 ± 19.24 | 312.50 ± 75.54 | 254.25 ± 93.27 | 0.39 |

Data are presented as mean ± SD. One-way ANOVA test was conducted for calculating statistical difference.
